# Ketogenic diet promotes tumor ferroptosis but induces relative corticosterone deficiency that accelerates cachexia

**DOI:** 10.1101/2023.02.17.528937

**Authors:** Miriam Ferrer, Nicholas Mourikis, Emma E. Davidson, Sam O. Kleeman, Marta Zaccaria, Jill Habel, Rachel Rubino, Thomas R. Flint, Claire M. Connell, Michael J. Lukey, Eileen P. White, Anthony P. Coll, Ashok R. Venkitaraman, Tobias Janowitz

## Abstract

The dependency of cancer cells on glucose can be targeted with high-fat low-carbohydrate ketogenic diet (KD). However, hepatic ketogenesis is suppressed in IL-6 producing cancers, which prevents the utilization of this nutrient source as energy for the organism. In two IL-6 associated murine models of cancer cachexia we describe delayed tumor growth but accelerated onset of cancer cachexia and shortened survival when mice are fed KD. Mechanistically, we find this uncoupling is a consequence of the biochemical interaction of two simultaneously occurring NADPH-dependent pathways. Within the tumor, increased production of lipid peroxidation products (LPPs) and, consequently, saturation of the glutathione (GSH) system leads to ferroptotic death of cancer cells. Systemically, redox imbalance and NADPH depletion impairs the biosynthesis of corticosterone, the main regulator of metabolic stress, in the adrenal glands. Administration of dexamethasone, a potent glucocorticoid, improves food intake, normalizes glucose homeostasis and utilization of nutritional substrates, delays onset of cancer cachexia and extends survival of tumor-bearing mice fed KD, while preserving reduced tumor growth. Our study highlights that the outcome of systemic interventions cannot necessarily be extrapolated from the effect on the tumor alone, but that they have to be investigated for anti­cancer and host effects. These findings may be relevant to clinical research efforts that investigate nutritional interventions such as KD in patients with cancer.

**Graphical abstract:** 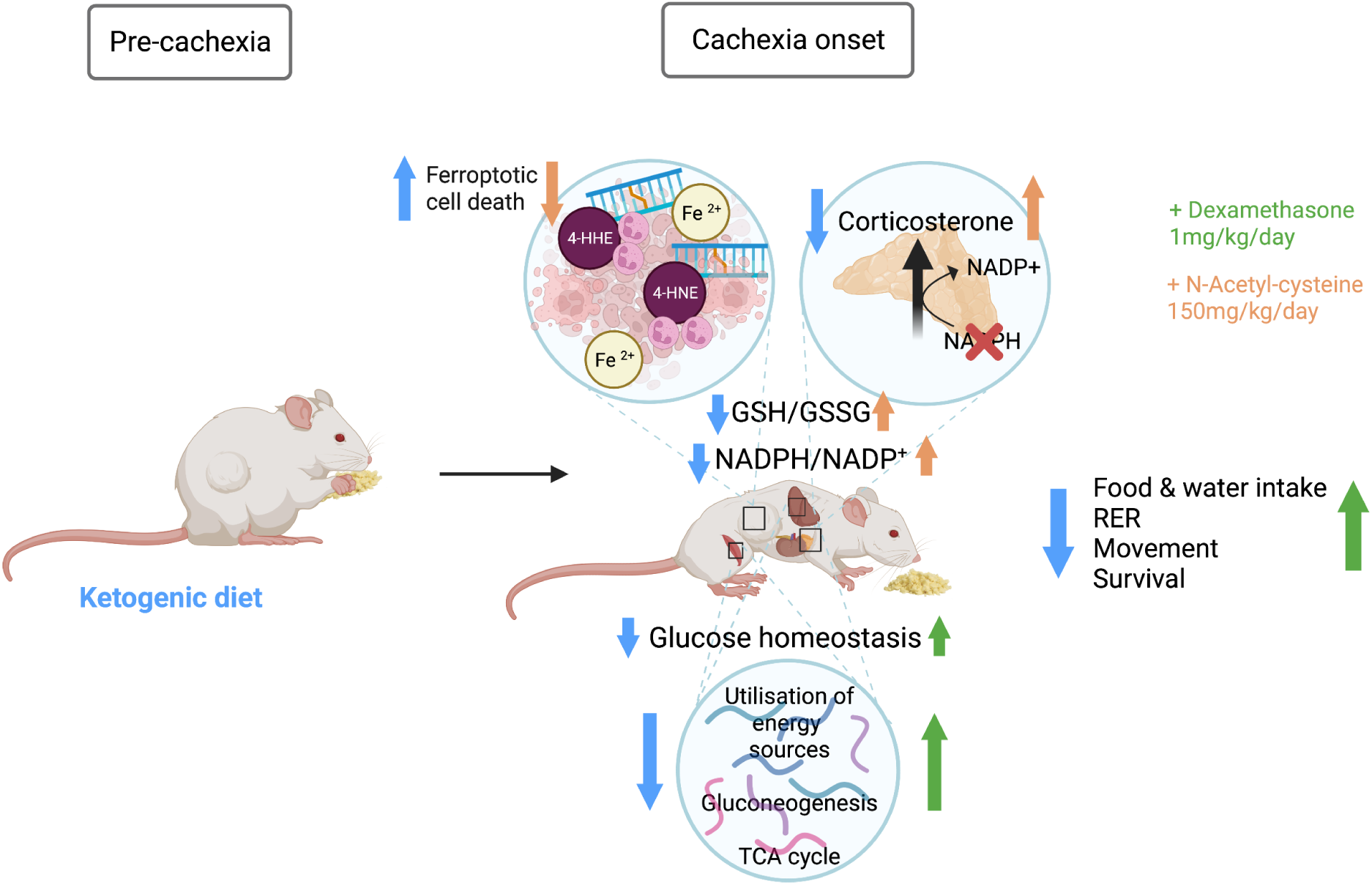

**HIGHLIGHTS:**

- Ketogenic diet delays tumor growth but accelerates cancer cachexia and shortens survival
- In the tumor, accumulation of lipid peroxidation products results in saturation of the GSH detoxifying pathway and ferroptotic death of cancer cells
- In the host organism, systemic redox state imbalance causes NADPH depletion, GDF-15 elevations, and relative corticosterone deficiency
- Dexamethasone coadministration with ketogenic diet delays onset of cancer cachexia by improving food intake, glucose homeostasis and utilization of nutritional substrates

## INTRODUCTION

Cancer, at a cellular and organismal level, is at least in part a metabolic disease. Cancer cells themselves have altered metabolism to accommodate nutrient demand and maintain growth and proliferation. For example, they frequently rely on increased glucose consumption to supply anabolic metabolism (Warburg 1925). At a systemic level, cancer can also alter the metabolism of the host by inducing profound changes in nutrient intake and handling that culminate in cachexia. Cancer cachexia is a severe wasting syndrome characterized by reduced food intake and terminal weight loss that affects up to 80% of all patients with cancer (von Haehling and Anker 2014), and causes significant morbidity and mortality (Farkas et al. 2013). The involuntary weight loss suffered by patients with cancer cachexia cannot be reversed by nutritional supplementation (Fearon et al. 2011), and this persistent metabolic stress condition increases glucocorticoids levels in humans and mouse models of cancer cachexia.

The dependency of cancer cells on glucose has been targeted by utilization of ketogenic diets (KD) containing high levels of fats and low levels of carbohydrates. KDs are explored as therapeutic intervention in end-stages of cancer that are associated with cachexia (Jansen and Walach 2016). Several studies report an anti-inflammatory and a delayed tumor growth effect of KD in pre-clinical models (Nakamura et al. 2018; Otto et al. 2008; Seyfried et al. 2003) and in humans (Schwartz et al. 2015). However, hepatic ketogenesis is suppressed in murine models of cancer progression that are associated with interleukin-6 (IL-6) elevation and ultimately cause cancer cachexia (Flint et al. 2016; Goncalves et al. 2018). This results in an inability of the organism to convert KD to survival-sustaining energy supply molecules. Together this raises the question whether the inability of the cancer cell or the organism to utilize the macronutrients supplied by KD is dominant with regard to outcome.

Lipid peroxidation is a non-enzymatic route of fatty acid metabolism that is oxygen radical-dependent (Massey and Nicolaou 2011; Yin, Xu, and Porter 2011). Metabolism of fats through non-enzymatic lipid peroxidation is a recognized source of highly reactive molecules named lipid peroxidation products (LPPs), such as 4-hydroxynonenal (4-HNE) and malondialdehyde (MDA), that cause cross-linkage on DNA and proteins through the formation of etheno-adducts (Goldstein 1975; Nair et al. 2019). Under physiological conditions, lipid peroxidation rates are low and non-toxic since LPPs are quickly removed from cells by constitutive antioxidants defense systems such as the NADPH-dependent glutathione (GSH) system (Little and O’Brien 1968; Ursini et al. 1982). When LPP detoxification fails, the accumulation of lipid peroxides results in a type of programmed cell death dependent on iron that is termed ferroptosis (Yang et al. 2014).

Cortisol is the major human glucocorticoid, equivalent to corticosterone in rodents. Cortisol release is part of the physiological response to starvation in cancer cachexia that drives adaptive pathways and regulates nutrient storage and processing, glucose levels, protein breakdown (Hoberman 1950; Wing and Goldberg 1993), and lipolysis (Divertie, Jensen, and Miles 1991). Biosynthesis of glucocorticoids occurs in the cortex of the adrenal gland through repeated NADPH-dependent enzymatic reduction of cholesterol. This process is under the control of the hypothalamic-pituitary-adrenal (HPA) axis. The inability to mount an adequate stress response due to irreversible damage to the adrenal cortex (e.g., auto-immunity) (Bratland et al. 2009) or pharmacotherapy-induced suppression of the HPA axis (Henzen et al. 2000) presents a life-threatening condition.

Therefore, glucocorticoid synthesis and the pathway of LPP detoxification share the requirement for NADPH as cofactor, yet their biochemical interdependency has not been explored. This interaction becomes relevant when both pathways simultaneously occur in metabolically stressed organisms (e.g., cachexia) fed a diet with high fat content (e.g., ketogenic diet).

In this study, we set out to determine the differential effect of KD on tumors and the host organism using two murine models of cancer cachexia. We find that although KD slows tumor growth, it shortens survival by accelerating the onset of cachexia. Mechanistically, increased production of LPPs in KD-fed tumor-bearing mice leads to a systemic redox state imbalance. Within tumors, this results in saturation of the GSH detoxifying pathway and consequent ferroptotic death of cancer cells. Moreover, we discover that NADPH depletion impairs corticosterone biosynthesis in the adrenal cortex, inducing a relative hypocorticosteronemia and metabolic maladaptation in mice fed KD. Treatment with the synthetic corticosteroid dexamethasone delays the onset of cancer cachexia and extends survival of tumor-bearing mice fed KD by improving food intake, metabolic homeostasis and utilization of nutritional substrates while preserving reduced tumor growth.

The uncoupling of tumor growth from overall survival illustrates why clinical trials should monitor the host response to nutritional interventions such as KD closely.

## RESULTS

### Ketogenic diet delays tumor growth but shortens overall survival in two mouse models of cancer cachexia

To investigate the effect of ketogenic diet on established IL-6-secreting cachexia-inducing cancers and the tumor-bearing host, BALB/c mice bearing subcutaneous C26 colorectal tumors for 14 days, and KPC mice (Kras^G12D/+^;Trp53^R172H/+^;Pdx-1-Cre), a genetically engineered mouse model (GEMM) of pancreatic cancer, with >3-5mm size tumors were challenged with a low-carbohydrate, moderate-protein, high-fat diet (KD) or maintained on normal diet feeding (NF) (Figure S1A and Table S1). Tumor growth was significantly decelerated in mice fed KD in both models, indicating a diet-mediated anti-tumor effect (Figures 1A and 1B). However, KD prompted an earlier onset of cancer cachexia (>15% bodyweight loss) and systemic wasting, thus shortening overall survival (OS) in both C26 and KPC mice fed KD compared to their counterparts fed NF (Median OS: 10 days C26/KD, 14 days C26/NF, 6.5 days KPC/KD, 17 days KPC/NF) (Figures 1C and ID, Figures SIB and SIC). At endpoint, tumor-bearing mice exhibited loss of subcutaneous and gonadal fat tissue, depletion of quadriceps muscle mass and splenomegaly, recognized signs of cachexia (Figures SID and S1E).

**Figure 1.**
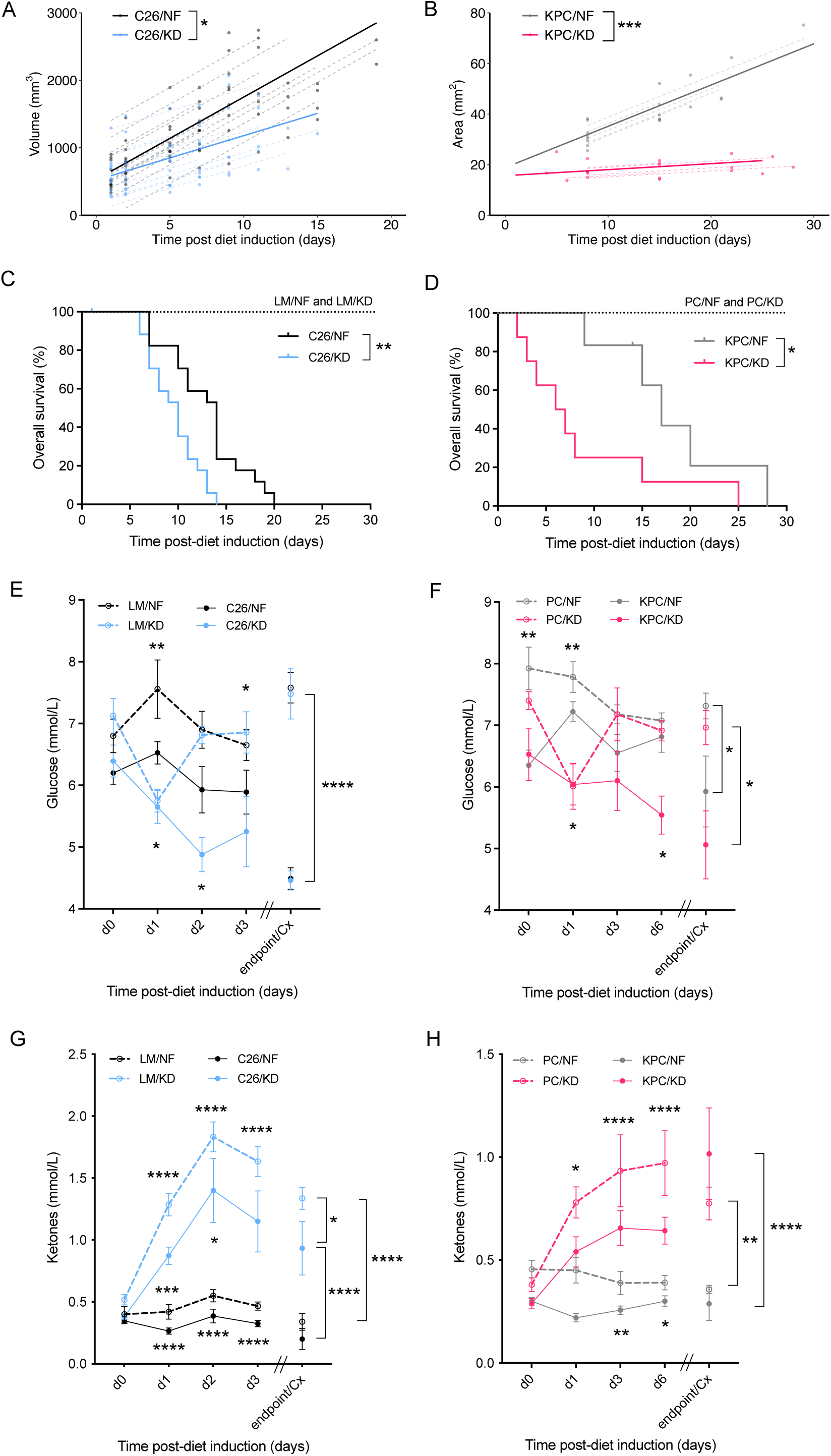
Ketogenic diet slows down tumor growth but shortens overall survival in C26 and KPC murine models of cancer cachexia. (A) Longitudinal tumor volume in C26 tumor-bearing mice fed ketogenic (KD) or standard (NF) diets (n=12). (B) Longitudinal tumor area in KPC tumor-bearing mice fed KD or NF (n=8). (C) Overall survival of C26 tumor-bearing mice and littermate controls on KD or NF (n=7 LM, n=17-18 C26). (D) Overall survival of KPC tumor­bearing mice and PC controls fed KD or NF (n=6-8). (E-F) Longitudinal glucose measurements from day 0 of diet change to cachectic endpoint in C26 tumor-bearing mice and littermate controls (n=5 LM, n=20 C26) (E), and in KPC tumor-bearing mice and PC controls (n=10) (F), fed either KD or NF diets. (G-H) Longitudinal ketone measurements from day 0 of diet change of cachectic endpoint in C26 tumor-bearing mice and littermate controls (n=5 LM, n=20 C26) (G), and in KPC tumor-bearing mice and PC controls (n=10) (H), fed either KD or NF diets. Data are expressed as the mean ± SEM. Overall survival (OS): time until mice reach >15% bodyweight loss. Differences in (A-B) were assessed by fitting a mixed effect model with coefficients for the intercept, slope and the difference in the slope between diets, and a random component for each individual mouse. Kaplan-Meier curves in (C-D) were statistically analyzed by using the log-rank (Mantel-Cox) test. Two-way ANOVA statistical tests with Tukey’s correction for post hoc comparisons were performed in (E-H). * p-value < 0.05, ** p-value < 0.01, *** p-value < 0.001, **** p-value < 0.0001.

Longitudinal monitoring of blood glucose levels in mice bearing established tumors showed an acute decrease in glucose of littermate controls (LM) and PC controls after introduction of KD that completely recovered after 24h to similar levels of controls fed NF. In contrast, C26 and KPC tumor-bearing mice on KD did not adapt to the new nutritional source and their glucose levels kept declining over time, reaching the lowest levels at cachectic endpoint. Tumor-bearing mice on NF had lower glucose levels than their non-tumor bearing counterparts but higher than tumor-bearing mice on KD, and they were able to maintain stable glucose measurements until onset of cachexia when the levels dropped abruptly (Figures IE and IF). Circulating ketones were significantly increased in all KD fed compared to NF fed groups, but tumor-bearing mice had lower levels compared to their non-tumor bearing control counterparts on the same diet (Figures 1G and 1H). Littermate mice on KD had a 2-fold upregulation of hepatic mRNA levels of PPARα target genes that regulate ketogenesis, *Acadm* and *Hmgsc2,* compared to littermates on NF. The expression was also 2-fold higher than in C26 tumor-bearers on KD, which were unable to upregulate the transcriptional targets responsible for ketogenesis despite the increased dietary substrate. C26 tumor-bearing mice on NF exhibited suppressed transcriptional regulation of ketogenesis compared to NF fed littermates, as previously described (Flint et al. 2016) (Figures S1F and S1G). Food intake was decreased in a similar manner in both tumor-bearing groups as cachexia developed (Figures S1H and S1I). Taken together, these data demonstrate not only that KD does not overcome the tumor-mediated metabolic reprogramming of the liver that suppresses ketogenesis, but it also impairs metabolic responses that maintain glucose homeostasis in the tumor-bearing host.

### Ketogenic diet induces formation of etheno-adducts and ferroptotic cell death of cancer cells that can be prevented by NAC

Detoxification of reactive LPPs by the constitutive antioxidant glutathione (GSH) is a NADPH-demanding process (Figure S2A)(Pannala et al. 2013). To quantify the formation of lipid peroxidation products in the context of the lipid-enriched KD (Table S1) and investigate the redox state of both the tumor and the host, we performed an in-depth metabolomics analysis on liver and tumor tissue of tumor-bearing C26 mice and control mice fed KD or NF. 4-HNE, the major lipid peroxide resulting from oxidation of fatty acids, accumulated in the liver of tumor-bearing C26 mice fed KD compared to NF fed tumor-bearing mice. Littermates without tumors on KD had unchanged levels of 4-HNE in the liver compared to NF fed littermates, suggesting its production was efficiently detoxified (Figure 2A). GSH to GSSG ratio in the liver was decreased in C26 tumor-bearing mice on KD compared to those on NF, suggesting an ongoing utilization of the reductive power of glutathione molecules with the purpose of detoxifying LPPs (Figure 2B). Tumor metabolomics of C26 mice fed KD or NF were separated by PCA (Figure S2B). GSH to GSSG ratio in tumors from C26 mice on KD was lower indicating a diminished reductive potential, in keeping with GSH consumption for the detoxification of LPPs (Figure 2C). The rate-limiting precursor metabolite for GSH biosynthesis, cysteine, was also decreased in tumors from C26 mice fed KD (Figure 2D), whereas ophthalmate, a biomarker for oxidative stress and GSH depletion (Dello et al. 2013), significantly accumulated in KD tumors (Figure S2C). Other indicators of redox perturbation, such as collapse of the antioxidant carnosine (Scuto et al. 2020) (Figure S2D) and evidence of hypotaurine to taurine oxidation (Figures S2E and S2F) (Fontana et al. 2005) were present in tumors of C26 KD-fed mice. These data are compatible with an increased formation of toxic and highly mutagenic lipid peroxides that saturates the GSH pathway in tumors from C26 mice on KD compared to those fed NF. Adduct formation by the lipid peroxide 4-HNE was significantly elevated in the tumors of C26 and KPC tumor-bearing mice fed KD compared to those fed NF (Figure 2E and 2F). 4-HNE adducts in KD-fed C26 mice were prevented with administration of N-acetyl cysteine (NAC), an antioxidant that boosts the GSH pathway by increasing GSH biosynthesis and therefore clearance of LPPs (Figures 2E, S2A and S2G).

**Figure 2.**
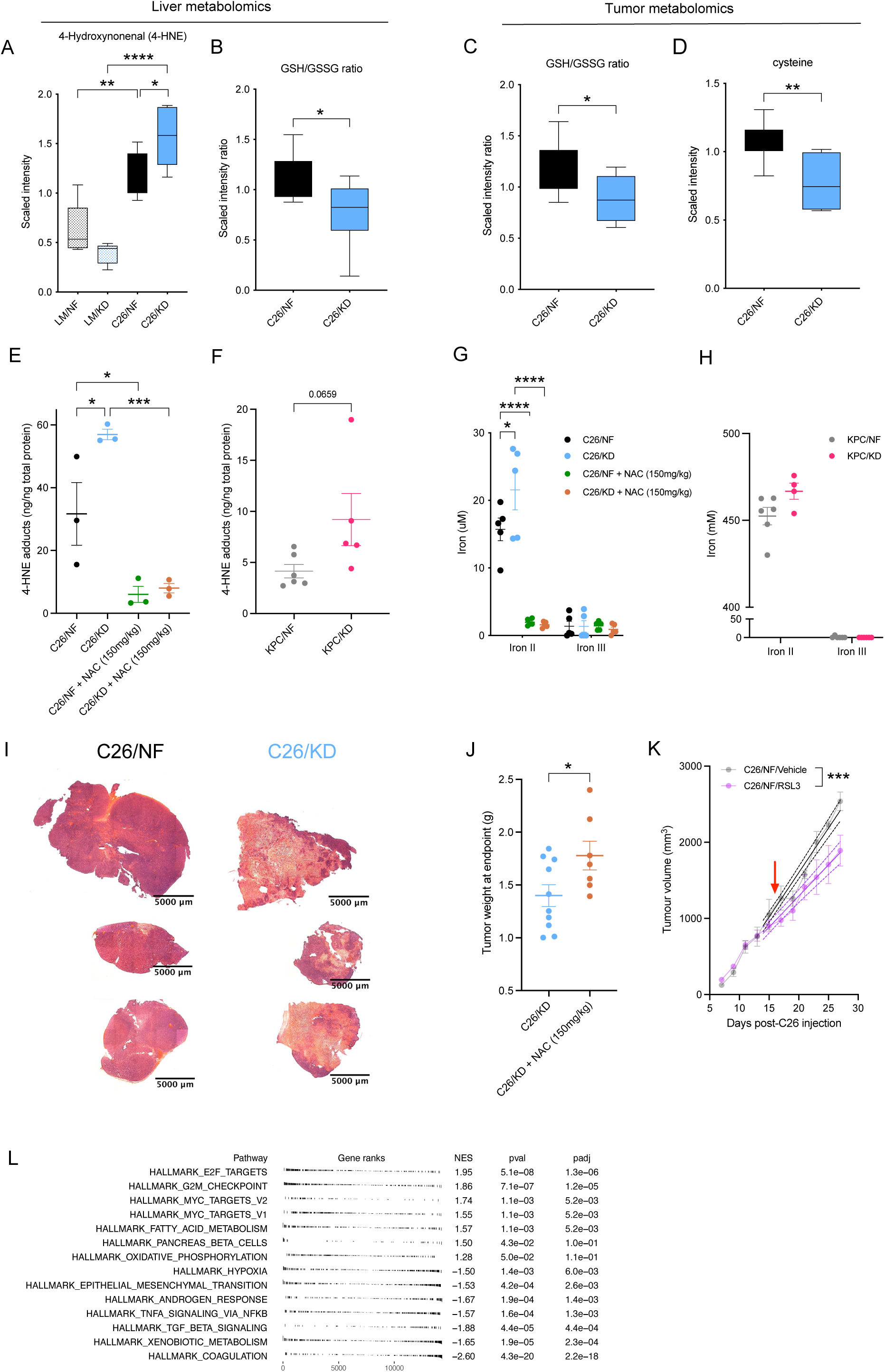
Ketogenic diet induces ferroptotic cell death of cancer cells that can be prevented by NAC. (A-B) Quantification by UPLC-MS/MS of 4-hydroxynonenal (4-HNE) and GSH/GSSG ratio (B) in the liver of C26-tumor bearing and littermate controls on KD or NF diets (n=7). (C-D) Quantification by UPLC-MS/MS of GSH/GSSG ratio (C) and cysteine (D) in the tumor of C26 mice (n=7). (E) Detection of 4-HNE adducts in tumor lysates from C26 tumor-bearing mice untreated or treated with N-acetyl cysteine (NAC) and fed KD or NF (n=3). (F) 4-HNE adducts formation in tumor of KPC mice fed KD or NF (n=4-6). (G-H) Concentration of ferric (III) and ferrous (II) iron in the tumor of C26 mice fed KD or NF, untreated or treated with NAC intraperitoneally (n=5) (G), and in tumors of KPC mice fed either diet (n=6) (H). (I) Haematoxylin and eosin (H&E) staining of tumors from C26 mice fed KD or NF. (J) Weight of tumors at the time of cachexia in C26 tumor­bearing mice fed KD untreated or treated with NAC (n=7-10). (K) Longitudinal tumor volume of C26 mice fed standard diet and treated with RSL3 or vehicle control (n=4-6). (L) GSEA of upregulated and downregulated pathways in tumors of KPC mice fed KD compared to KPC fed NF (n=5). Data are expressed as the mean ± SEM. One-way ANOVA with Tukey’s correction for post hoc testing was used in (A, E-G). Statistical differences in (B-D, H-I) were examined using an unpaired two-tailed Student’s t-test with Welch’s correction. Simple linear regression model was applied to (J). Statistical analysis in (K) is described in Methods. * p-value < 0.05, ** p-value < 0.01, *** p-value < 0.001, **** p-value < 0.0001.

Accumulation of lipid peroxides results in a type of programmed cell death dependent on iron named ferroptosis (Dixon et al. 2012) that has previously been described in cysteine-depleted tumors (Badgley et al. 2020). To explore the possibility that ferroptotic cell death in tumors from KD-fed mice is partly responsible for reduced tumor burden and slower tumor growth trajectories, we first performed Oil-Red-O staining of lipids in tumors from C26 mice. C26 mice on KD had significant accumulation of lipid droplets in the tumor tissue, whereas tumors from C26 mice on NF had no evidence of lipid storage (Figure S2H). We next quantified the levels of ferrous (Fe^2+^/Iron II) and ferric (Fe^3+^/ Iron III) iron in tumor samples. Ferrous iron is an established indirect marker of ferroptosis and it accumulated in tumor tissues from C26 and KPC mice fed KD, suggesting an ongoing ferroptotic cell death mechanism in these tumors compared to tumors from mice fed NF (Figure 2G and 2H). A dead tumor core was observed in tumors from mice fed KD by hematoxylin and eosin (H&E) staining (Figure 21), supporting an ongoing cell death process in these tumors. Moreover, tumors of mice on KD treated with NAC were notably bigger than those in untreated mice fed KD (Figure 2J), and ferrous iron levels were depleted upon NAC treatment (Figure 2G), indicating that NAC administration is sufficient to prevent ferroptosis and cell death in these tumors. We performed a pharmacological experiment to support this hypothesis, with the aim to recapitulate the effect induced by KD. Indeed, administration of RSL3, a specific inhibitor of glutathione peroxidase 4 (GPx4) that blocks the detoxification of LPPs by the GSH pathway and thereby induces cellular ferroptosis, to tumor-bearing mice fed NF led to reduced tumor growth (Figure 2K).

We next set out to validate these findings in the KPC model. RNAseq data from tumors of cachectic KPC mice demonstrated significant overexpression of E2F and Myc targets, which are major regulators of cellular metabolism in response to stress (Dong et al. 2020), in tumors of KPC mice fed KD compared to KPC fed NF. The E2F axis has previously been associated to a role as promoter of oxidative stress and ferroptosis in neurons, and its silencing leads to prevention of this iron-dependent cell death (Mishima 2021). Myc signalling has also been linked to mediation of, and sensitization to, ferroptotic cell death (Alborzinia et al. 2022; Lu et al. 2021). Gene expression marking activity of the G2/M checkpoint were significantly upregulated in tumors of tumor-bearing KPC mice fed KD, and since the activation of this checkpoint prevents cells from entering mitosis when the DNA is damaged and allows for its repair, this suggests that LPP formation and reactivity in the cancer cells causes DNA crosslinking, adduct formation and consequently cell cycle arrest. The activity of the cytochrome P450 reductase (POR), which has a major role in the metabolism of drugs and xenobiotics, requires NADPH (Esteves, Rueff, and Kranendonk 2021). Downregulation of pathways related to xenobiotic metabolism in the tumors of mice fed ketogenic diet can be explained by the ongoing systemic NADPH depletion in these mice (Figure 2L).

Apoptosis was disregarded as an additional mechanism contributing to reduced tumor growth in mice fed KD because apoptotic markers such as Caspase-3 were expressed similarly in Western Blot (WB) of C26 tumors from both dietary groups. BAX staining suggested even lower levels of apoptotic cell death in tumors from C26 KD fed mice compared to those from C26 fed NF (Figure S2I). Cell proliferation, measured by Kİ67 staining, was also not affected by the dietary challenge itself (Figure S2J).

All of these findings demonstrate that elevated oxidative stress leading to cell cycle arrest and ferroptotic cell death caused by build-up of LPPs are mechanisms contributing to smaller tumor burden in mice fed KD.

Since ferroptosis is an immunogenic process (Efimova et al. 2020), we next studied the immune infiltration in tumors from KD- and NF-fed mice. We found a positive trend for enrichment of all immune cell types examined in KD tumors, reaching statistical significance for neutrophils (Figures S2K, S2L, S2M and S2N). This observation may be explained by active recruitment of neutrophils to areas undergoing ferroptotic cell death.

### Ketogenic diet impairs glucocorticoid synthesis in tumor-bearing mice

Corticosterone is the main glucocorticoid involved in metabolic adaptation under stress conditions in mice that acts as a regulator of metabolic rates and availability of fuel substrates. Metabolic stressors such as caloric restriction associated with cachexia induce high levels of corticosterone (Flint et al. 2016). Similar to the detoxification of LPPs, the corticosteroid synthesis pathway in the cortex of the adrenal gland requires a constant supply of NADPH cofactor molecules (Figure S3A). To gain a deeper understanding of the metabolic stress response in KD and NF fed mice, we quantified circulating levels of corticosterone and cholesterol, the substrate for corticosterone biosynthesis in the adrenals. Control littermates on either diet had baseline normal levels of corticosterone in circulation. At the time of cachexia, tumor-bearing C26 mice fed NF displayed a sharp increase in corticosterone concentration, but levels in tumor-bearing C26 mice fed KD were not elevated in comparison. (Figures 3A and S3B). Cholesterol availability was similar in C26 and KPC mice fed with KD and NF diets (Figures 3B and 3C) but pregnenolone levels significantly accumulated in the plasma of C26 mice fed KD compared to those fed NF (Figure 3D), pointing towards a defect in the synthetic cascade of corticosterone (Figure S3A) rather than an absence of substrate.

**Figure 3.**
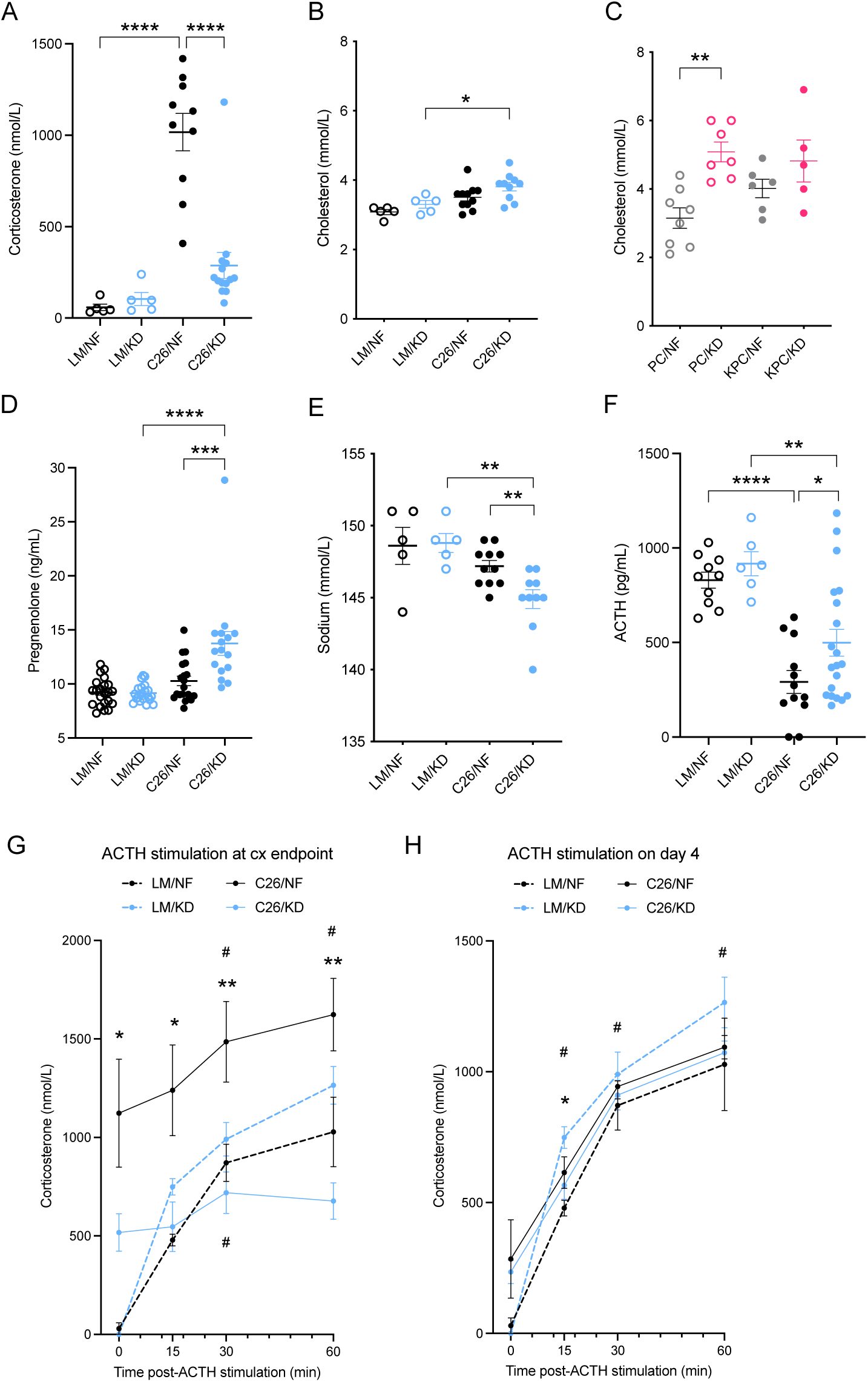
Ketogenic diet induces relative corticosterone deficiency in C26-tumor bearing mice. (A) Corticosterone hormone levels in plasma of cachectic C26 tumor-bearing mice and littermate controls fed KD or NF diets (n=5 LM, n=10-14 C26). (B-C) Plasma cholesterol levels in cachectic C26-tumor bearing mice and littermate controls (n=5 LM, n=10-ll C26) (B), and in cachectic KPC tumor-bearing mice and PC controls (n=5-8) (C) fed KD or NF diet. (D) Pregnenolone hormone levels in plasma of cachectic C26 tumor-bearing mice and littermate controls on KD or NF diets (n=16-22). (E) Sodium levels in plasma of cachectic C26-tumor bearing mice and littermate controls on KD or NF diets (n=5 LM, n=10-ll C26). (F) Levels of adrenocorticotropic hormone (ACTH) in plasma of cachectic C26 tumor-bearing mice and littermate controls fed KD or NF (n=6-10 LM, n=12-20 C26). (G-H) Synacthen test and quantification of corticosterone response at baseline and 15, 30 and 60 minutes after ACTH stimulation in cachectic C26 tumor-bearing mice and littermate controls (endpoint) (G), or only 4 days after diet change (18 days after C26 cell injection) (n=5) (H). Data are expressed as the mean ± SEM. One-way ANOVA with Tukey’s correction for post hoc testing was used in (A-D, F). Two-way ANOVA statistical tests with Tukey’s correction for post hoc comparisons were performed in (G-H). * p-value < 0.05, ** p-value < 0.01, *** p-value < 0.001, **** p-value < 0.0001, # p-value < 0.05 compared to time = 0.

We next compared the transcriptome of the adrenal glands in tumor-bearing KPC mice fed KD or NF, and using Gene Set Enrichment Analysis (GSEA) we found that the steroid biosynthesis and cholesterol homeostasis pathways were significantly downregulated in KPC mice fed KD compared to those fed NF (Figure S3C). These data demonstrate an impaired corticosterone production and inefficient stress response in the adrenal glands of tumor­bearing mice fed KD compared to those fed NF. One of the major actions of aldosterone, a mineralocorticoid hormone derived from downstream processing of corticosterone in the adrenal glands, is sodium retention and potassium loss. Quantification of circulating sodium in C26 mice and controls revealed a relative hyponatremia in C26 tumor-bearing mice fed KD compared to C26 fed NF and littermate controls (Figures 3E), further supporting impaired hormone biosynthesis in the adrenal glands of these mice.

The adrenal glands are part of the HPA axis, which regulates a cascade of endocrine pathways including the production of corticosterone (cortisol in humans). The adrenocorticotropic hormone (ACTH) is a hormone produced by the pituitary gland that binds its receptor in the cells of the zona fasciculata of the adrenal cortex and drives the production of corticosterone. Elevated circulating corticosterone levels induce negative feedback on the hypothalamus and inhibit ACTH release. In order to assess whether the impaired synthesis of corticosterone in tumor-bearing mice fed KD is a) a localized phenomenon in the cortex of the adrenal glands, b) due to an upstream defect in the HPA axis, such as inadequate ACTH production by the pituitary gland, or c) a combination of both, we quantified ACTH in the plasma of cachectic mice and littermate controls fed NF or KD. At endpoint, cachectic tumor­bearing mice fed KD showed significantly higher levels of plasma ACTH compared to tumor­bearing mice fed NF, supporting the hypothesis of an intrinsic deficiency in corticosterone biosynthesis in the adrenal glands (Figure 3F). However, given the variance in ACTH levels, a minor contribution from upstream mechanisms to the observed hypocorticosteronemia cannot be ruled out.

Since corticosterone release can potentially be driven by direct stimulation of the adrenal glands by non-ACTH peptides such as IL-6 (Bethin, Vogt, and Muglia 2000; Salas et al. 1990; Žarković et al. 2008), and the C26 model is known to display high IL-6 levels (Flint et al. 2016), we quantified circulating levels of this cytokine. No diet-mediated differences between the tumor-bearing groups were observed (Figure S3D), indicating that IL-6 does not contribute to the relative corticosterone deficiency observed in KD-fed tumor-bearing mice.

To asses adrenal gland responsiveness at the onset of cachexia, we performed an ACTH stimulation experiment using synthetic ACTH (synacthen test). Baseline levels pre-stimulation were significantly elevated in tumor-bearing C26 mice on NF compared to tumor-bearing C26 mice on KD (Figure 3G). Upon ACTH injection, plasma corticosterone concentration increased over time in tumor-bearing C26 mice fed NF and littermates, while levels in tumor-bearing C26 mice fed KD did not significantly change. After 60 minutes, levels of corticosterone in tumor­bearing C26 mice fed NF were almost 2.5-fold higher than in tumor-bearing C26 mice fed KD. Both littermate groups showed similar responses and reached peak levels comparable to those of tumor-bearing C26 mice on NF at baseline (Figure 3G). These data point towards an intrinsic difficulty in the adrenal glands of KD-fed tumor-bearing mice to respond to hormonal stimulation and release corticosterone compared to NF-fed tumor-bearing mice. The same synacthen test was performed in a different mouse cohort four days after enrolment and diet change (daylδ post-C26 injection). Both tumor-bearing C26 groups had higher baseline corticosterone levels compared to the littermate controls. In response to ACTH administration, tumor-bearing C26 mice on NF had a stronger response to ACTH than their littermate controls. Conversely, corticosterone upregulation in tumor-bearing C26 mice on KD was significantly reduced compared to the response of control mice on the same diet (Figure 3H). Thus, our results indicate early signs of malfunction of the stress axis in tumor-bearing mice fed KD even only 4 days after dietary change, but these only become evident and clinically relevant at the onset of cachexia, when an adequate stress response that coordinates adaptation and systemic homeostasis is essential. Altogether these data provide evidence that KD drives the development of a relative hypocorticosteronemia in tumor-bearing C26 mice fed KD compared to those fed NF.

### NAC treatment rescues corticosterone synthesis in tumor-bearing mice fed ketogenic diet

In order to identify the mechanism underlying the relative deficiency in corticosterone biosynthesis in tumor-bearing mice fed KD, we next explored the interaction of the GSH pathway (Figure S2A) and the corticosterone synthesis pathway (Figure S3A) through their common need of NADPH sources. Targeted quantification of NADPH in the adrenal glands of cachectic mice and controls showed increased levels of NADPH in tumor-bearing C26 mice fed NF, as it would be anticipated in the context of an ongoing release of corticosterone. However, the NADPH supply was diminished in the adrenal glands of tumor-bearing C26 mice fed KD. Administration of NAC, a cysteine prodrug that replenishes intracellular GSH levels in the absence of NADPH consumption, rescued NADPH levels in these mice (Figure 4A). Taken together, these findings indicate that the increased demand for NADPH in the process of detoxification of LPPs leads to a shortage of this cofactor molecule, which then is not available for use in the synthesis of corticosterone and leads to low levels of this stress hormone in tumor-bearing mice fed KD.

**Figure 4.**
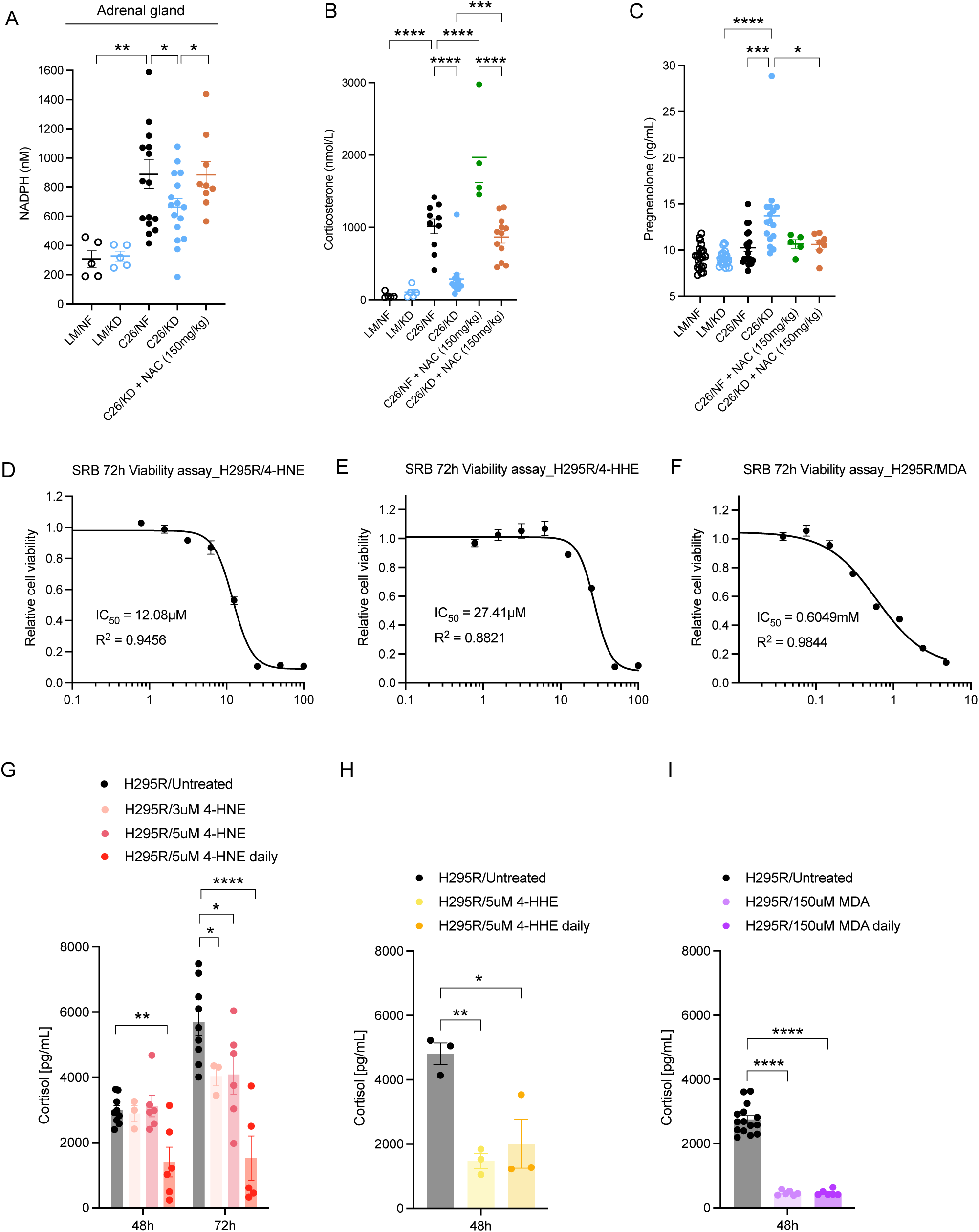
Effect of lipid peroxidation products (LPPs) on adrenal function and rescue with NAC. (A) NADPH quantification in the adrenal glands of littermate controls and C26 tumor-bearing mice fed KD or NF, and C26 tumor-bearing mice fed KD treated with NAC (n=5 LM, n=9-ll C26). (B-C) Corticosterone (n=5 LM, n=10-14 C26) (B) and pregnenolone (n=5-22) (C) levels in plasma of littermate controls and C26 tumor-bearing mice fed KD or NF, untreated or treated with NAC. (D-F) SRB assays of H295R cells treated for 72h with increasing concentrations of 4-HNE (D), 4-HHE (E) or MDA (F) (n=3 independent experiments). Viability is expressed relative to vehicle-treated control cells. (G-I) Cortisol levels in the media of H295R cells treated with 4-HNE (n=3-6) (G), 4-HHE (n=3-6) and MDA (n=6-15) (I) at 48h and 72h timepoints. Data are expressed as the mean ± SEM. One-way ANOVA with Tukey’s correction for post hoc testing was used in (A-C, G-l). * p-value < 0.05, *** p-value < 0.001, **** p-value < 0.0001.

To examine this hypothesis further, we measured corticosterone levels in tumor-bearing mice fed KD or NF and treated with NAC. Circulating corticosterone was markedly higher in the NAC-treated groups compared to untreated and control groups on the same diet (Figure 4B). Simultaneously, pregnenolone accumulation in KD-fed tumor-bearing mice was no longer detected upon NAC treatment (Figure 4C) indicating conversion of this early intermediate to downstream intermediates of corticosterone biosynthesis and ultimately to corticosterone. Therefore, promoting GSH production through NAC diminishes the need of NADPH oxidation, consequently increasing GSH’s LPP-detoxifying activity and at the same time preventing NADPH depletion. NADPH availability enables an appropriate synthesis of corticosterone in the adrenals and leads to the physiological rise in systemic corticosterone levels in the context of metabolic stress associated with cachexia.

### LPPs exposure decreases cortisol production in a human adrenal cortex-derived cell line

We next implemented the human adrenal cortex-derived cell line, H295R, to test the direct effects of LPPs on cortisol synthesis *in vitro. We* first identified for 4-hydroxynonenal (4- HNE), 4-hydroxyhexenal (4-HHE) and malondialdehyde (MDA) tolerated doses that did not affect cell viability as assessed by Sulforhodamine B (SRB) survival assays, a widely used method for *in vitro* cytotoxicity screening. After exposing the cell line to tolerated doses of 4-HNE (Figure 4D), we quantified the levels of cortisol released to the media at 48h and 72h after exposure to 4-HNE, in untreated cells, or in cells that were exposed to 5 µM 4-HNE once daily.

The results show significantly diminished cortisol concentration upon daily exposure to 5 µM 4- HNE, as well as reduced cortisol levels after 72h of exposing the cells to a single-dose of 3 µM or 5 µM 4-HNE (Figure 4G). Similarly, treatment of H295R cells with a tolerated dose of 4-HHE led to lower cortisol production compared to untreated adrenocortical cells (Figure 4E and 4H). This cortisol-suppressing effect was even more robust upon treatment of H295R cells with a tolerated dose of MDA (Figure 4F). Single exposure to 150 µM MDA led to a 6-fold decrease in cortisol production by H295R adrenocortical cells after 48h, and daily treatment did not amplify the inhibition compared to single treatment (Figure 41). Thus, these data suggests that LPPs exert a direct effect on cortisol production in adrenocortical human cells *in vitro*.

### GDF-15 is elevated in cachexia and increased by ketogenic diet

While the KD-mediated biochemical impairment of the adrenal glands stress response and resulting defective glucocorticoid biosynthesis in tumor-bearing mice described above can account for shortened survival, it does not necessarily explain reduction in food intake. GDF-15, a TGF-beta superfamily member that has been shown to induce reduced food intake by binding its cognate receptor GFRAL in the area postrema, is produced by cells under stress including metabolic stress (Patel et al. 2019). It has been implemented in the anorectic response in cancer cachexia (Hsu et al. 2017). In another model of aldehyde toxicity-induced anorexia, GDF- 15 reversibly modulated food intake (Mulderrig et al. 2021). In keeping with these findings, we observed elevated circulating GDF-15 levels in the C26 model system during cachexia in NF-fed mice, which were further elevated in KD-fed cachectic mice (Figure 5A), reflecting the systemic oxidative and metabolic stress of the organism and explaining at least in part the reduced food intake observed in the cachectic phase of the disease progression (Figure S1F and S1G).

**Figure 5.**
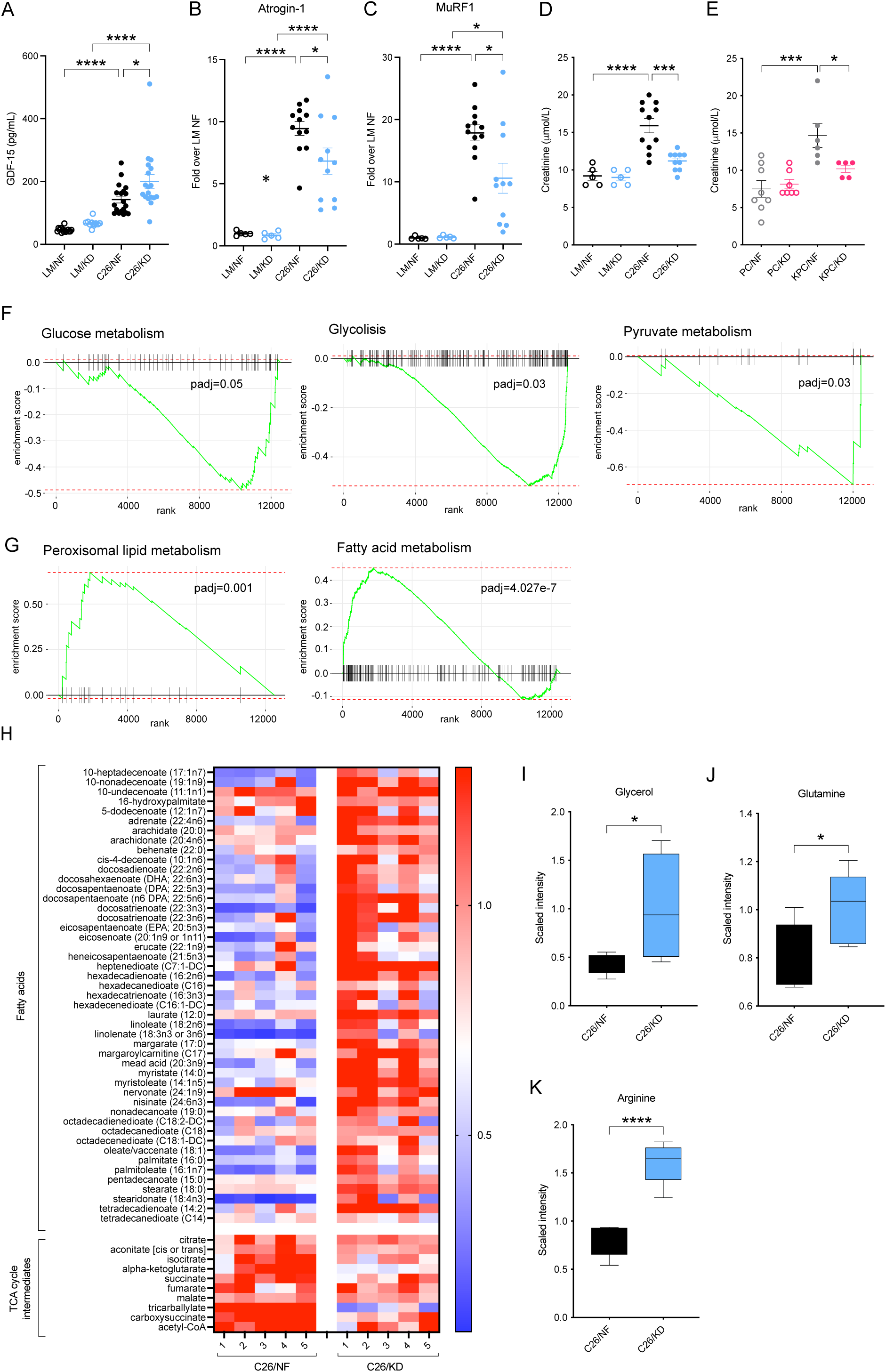
Appropriate usage of energy sources in the context of cachexia is impaired in KD-fed tumor-bearing mice. (A) Levels of GDF-15 in the plasma of C26 tumor-bearing mice and littermate controls fed KD or NF (n=ll-19). (B-C) mRNA levels of the E3 ligases Atrogin-1 (B) and MuRF1 (C) in the quadriceps of C26 tumor-bearing mice and littermate controls fed KD or NF (n=5 LM, n=12 C26). (D-E) Plasma creatinine levels in C26 tumor-bearing mice and littermate controls (n=5 LM, n=9-ll C26) (D), and KPC tumor-bearing mice and PC controls (n=5-8) (E) fed either KD or NF. (F-G) GSEA analysis of downregulated (F) and upregulated (G) pathways in KD-fed C26 tumor-bearing mice compared to those NF-fed (n=5). (H) Heatmap of metabolites and specific metabolic pathways in C26 tumor-bearing mice fed KD or NF (n=5). (I-K) Quantification by UPLC-MS/MS of the main substrates of the TCA cycle, glycerol (I), glutamine (J), and arginine (K) in the tumor of C26 mice fed KD or NF (n=7). Data are expressed as the mean ± SEM. One-way ANOVA with Tukey’s correction for post hoc testing was used in (A-E). Statistical analysis in (F-G) is described in Methods. Statistical differences in (I-K) were examined using an unpaired two-tailed Student’s t-test with Welch’s correction. * p-value < 0.05, *** p-value < 0.001, **** p-value < 0.0001.

### Appropriate usage of energy sources in the context of cachexia is impaired in ketogenic diet- fed tumor-bearing mice

To test the relevance of glucocorticoid-driven metabolic adaptation that promotes survival in the context of a cachectic tumor-bearing host and the impact that its malfunction may have in the context of KD feeding, we assessed and compared the systemic metabolic state of these mice.

At endpoint, tumor-bearing mice in both dietary groups exhibited signs of wasting and cachexia, including splenomegaly, and loss of fat and muscle mass (Figure S1D and S1E). Atrogin-1 and MuRF1 are markers of skeletal muscle atrophy and proteolysis of muscle proteins in various pathological conditions, including cachexia (Yuan et al. 2015). mRNA Quantification of these two muscle-specific E3 ubiquitin ligases in the quadriceps of cachectic mice exhibited upregulated expression in both tumor-bearing C26 groups compared to controls, yet the fold increase was significantly less pronounced in tumor-bearing KD fed than in NF fed mice (Figures 5B and 5C). Creatinine, an end product of muscle catabolism, was markedly increased in the circulation of cachectic tumor-bearing C26 and KPC mice fed NF but not in in cachectic tumor­bearing mice fed KD (Figures 5D and 5E), presumably as a consequence of lower corticosterone levels that regulate the breakdown of proteins. Urea production is commonly upregulated in cachexia due to increased protein release from muscle tissue breakdown (Corbello Pereira et al. 2004; Haines et al. 2019), however, urea levels stayed low in cachectic tumor-bearing mice fed KD compared to cachectic tumor-bearing mice on NF (Figures S4A and S4B). These observations indicate that, in cachectic tumor-bearing mice fed KD, the process of ubiquitin-mediated proteolysis is impaired despite exhibiting cachexia-associated muscle atrophy.

We next examined how the liver responded to the metabolic stress in cachectic tumor­bearing mice undergoing systemic wasting and decreased food intake. RNAseq data from livers of cachectic tumor-bearing KPC mice fed NF or KD demonstrated downregulated hepatic glycolysis, glucose metabolism and pyruvate metabolism in KD-fed KPC, all of which are pathways that are usually stimulated by glucocorticoids (Figure 5F). Peroxisomal lipid metabolism and fatty acid metabolism appeared upregulated in the liver of tumor-bearing KPC mice fed KD compared to those fed NF, in agreement with a predominantly lipid-rich diet (Figure 5G). Moreover, untargeted metabolomics of the liver of littermate controls and tumor­bearing C26 mice on either KD or NF diets manifested distinct hepatic metabolic profiles between the groups (Figure S4C). Specific pathway analysis led us to identify a marked accumulation of fatty acid metabolites in the liver of tumor-bearing C26 mice fed KD compared to those fed NF (Figure 5H). These data, together with the observation of macroscopically bigger liver sizes (Figure S1D), suggested that inappropriate macronutrient utilization of lipids could account for the inability to sustain glucose homeostasis and the build-up of unprocessed fat in the liver of tumor-bearing mice on KD. In addition, tricarboxylic acid (TCA) cycle metabolic intermediates appeared clearly upregulated in the liver of tumor-bearing C26 mice fed NF compared to C26 mice fed KD, which were unable to increase these metabolites to the same extent (Figure 5H). The Krebs or TCA cycle in hepatic cells is the metabolic progenitor pathway for gluconeogenesis and the main source of energy for the body (Anderson et al. 2018; Rui 2014). Glycerol and aminoacids, such as glutamine and arginine, are a major cellular carbon source for oxidative catabolism via the TCA cycle. These energy substrates were elevated in the liver of tumor-bearing C26 mice fed KD (Figure 51, 5J and 5K), reinforcing the idea of an impaired host metabolic activity and deficient energy production to sustain survival in these mice.

Altogether, these data provide evidence that the lack of an appropriate corticosterone release in tumor-bearing mice fed KD is associated with metabolic maladaptation, inability to use energy sources, failure to achieve glucose homeostasis, and ultimately earlier onset of cancer cachexia.

### Dexamethasone treatment extends survival and improves metabolic adaptation of tumor­bearing mice fed ketogenic diet

Given the relevance of the steroid hormone corticosterone in metabolic adaptation and efficiency of energy utilization under stress conditions such as cancer cachexia, we next tested the hypothesis that the survival disadvantage in tumor-bearing mice fed KD compared to NF fed is driven by the differences in corticosterone production.

BALB/c mice bearing subcutaneous C26 colorectal tumors for 14 days were either fed KD or NF and treated daily with 1mg/kg dose of Dexamethasone intraperitoneally. Dexamethasone is a synthetic long-acting potent glucocorticoid analogue, with a glucocorticoid activity of 30 relative to cortisol (Liu et al. 2013; Mager et al. 2003). Treatment with Dexamethasone extended survival in tumor-bearing C26 mice fed KD compared to all other tumor-bearing groups fed with either diet. Specifically, Dexamethasone led to a delay in the onset of cachexia and a major improvement in overall survival (OS) of C26 mice fed KD compared to the untreated group (Median OS: 10 days C26/KD, 33 days C26/KD + Dex). Dexamethasone treatment of NF fed tumor-bearing C26 mice also significantly extended overall survival (OS) (endpoint defined as >15% bodyweight loss) for 5 days (Median OS: 14 days C26/NF, 19 days C26/NF + Dex), but to a lower degree than on KD fed C26 mice (Figure 6A, Figure S5B). Moreover, Dexamethasone treatment prompted faster growth of tumors in C26 mice fed NF and therefore shortened progression free survival (PFS) (endpoint defined as tumor size > 2000 mm^3^) in these mice (Figures 6B and S5C). Tumor sizes in C26 mice fed KD were not significantly affected by Dexamethasone treatment and tumor weight remained unchanged, whereas tumors in Dex-treated C26 mice fed NF were increased almost 2-fold at endpoint (Figure 6C).

**Figure 6.**
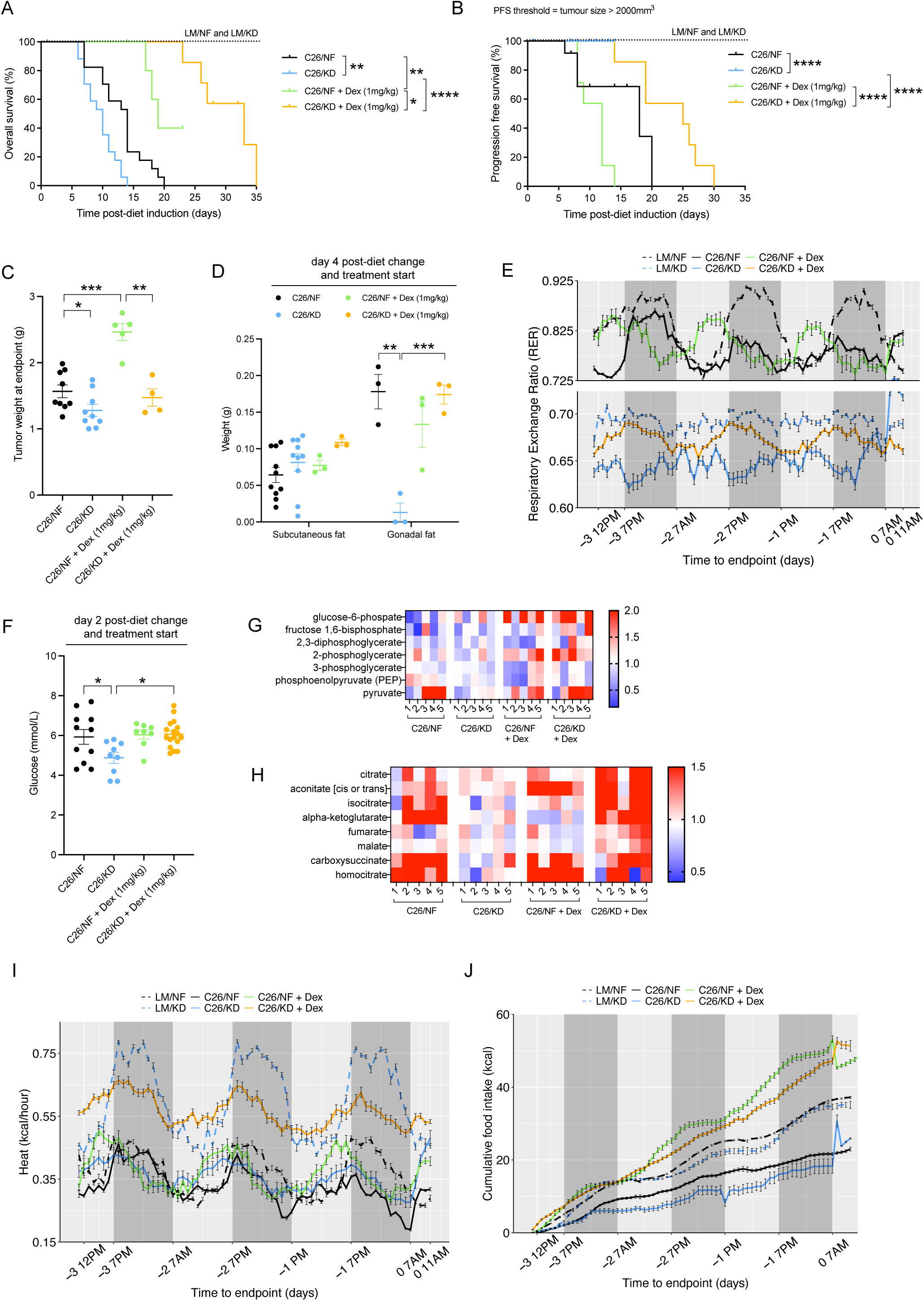
Dexamethasone treatment extends survival and improves metabolic adaptation of C26 mice fed ketogenic diet. (A-B) Overall survival (OS) (A) and Progression Free survival (PFS) (B) of C26 tumor-bearing mice fed KD or NF, untreated or treated with Dexamethasone, and littermate controls fed with either diet (n=7 LM, n=17-18 C26, n=7 C26 + Dex). (C) Weight of tumors in C26 tumor-bearing mice fed KD or NF, untreated or treated with Dexamethasone, at endpoint (n=9-10 C26, n=4-5 C26 + Dex). (D) Quantification of subcutaneous and gonadal fat stores in C26-tumor bearing mice fed KD or NF, untreated or treated with Dexamethasone, 4 days after diet change and start of treatment (n=3-10). (E) Cumulative food intake (kcal) during the last 4 days before endpoint in littermate controls and C26 tumor-bearing mice, untreated or treated with Dexamethasone, fed KD or NF (n=7). (F) Plasma glucose levels in C26-tumor bearing mice fed KD or NF, untreated or treated with Dexamethasone, 2 days after diet change and start of treatment. (G-H) Quantification by UPLC-MS/MS of metabolites involved in gluconeogenesis (G) and the TCA cycle (H) in the liver of C26 tumor-bearing mice on either KD or NF diets, untreated or treated with Dexamethasone (n=5). (I-J) Respiratory exchange ratio (RER) (I) and Heat (or energy expenditure) (J) during the last 4 days before endpoint in littermate controls and C26 tumor-bearing mice, untreated or treated with Dexamethasone, fed KDorNF. Data are expressed as the mean ± SEM. . Overall survival (OS): time until mice reach >15% bodyweight loss. Progression-Free Survival (PFS): time until tumor size reaches > 2000 mm^3^. Kaplan-Meier curves in (A-B) were statistically analyzed by using the log-rank (Mantel-Cox) test. One-way ANOVA with Tukey’s correction for post hoc testing was used in (C-D, F) Analysis in (E, G-J) is described in Methods. * p-value < 0.05,** p-value < 0.01, *** p-value < 0.001, **** p-value < 0.0001.

After 4 days of treatment with Dexamethasone, the depletion of fat stores induced by KD in tumor-bearing animals was rescued, but fat tissues in C26 mice fed NF showed no differences in weight upon Dexamethasone treatment (Figure 6D). This is in agreement with hepatic metabolomics data at endpoint, which shows decreased fatty acid metabolism in the liver of KD-fed C26 tumor-bearing mice treated with Dexamethasone compared to non-treated C26 tumor-bearing mice on KD. No changes in fatty acid metabolism were detected in NF-fed mice on Dexamethasone (Figure S5D). Quadriceps and liver mass were not affected by Dexamethasone in any of the groups, and splenic size was reduced in both C26 tumor-bearing groups, but more significantly in KD fed mice, as a consequence of the immunosuppressive effects of Dexamethasone (Figure S5E).

To investigate further the metabolic changes induced by corticosteroids during cancer progression and cancer cachexia in tumor-bearing mice on NF and KD diets, we next placed KD and NF fed tumor-bearing C26 mice, untreated or treated daily with 1mg/kg dose of Dexamethasone, and controls, in metabolic cages capable of precise monitoring of multiple metabolic parameters (details in Methods). The Respiratory Exchange Ratio (RER) is a ratio between the volume of CO_2_ being produced by the body and the amount of O_2_ being consumed. A value of RER close to 1.0 depicts the use of carbohydrates as a source of energy, whereas a ratio of 0.7 is indicative of fatty acids used as the primary fuel (Bhandarkar et al. 2021). RER values of littermate controls on NF fluctuated from carbohydrate-focused during night hours, when mice are physically more active and eating, to a more mixed energy source during light hours, when mice tend to sleep (Figure 6E). Littermate controls on KD exhibited moderate circadian rhythms in RER too, but values always depicted fat consumption as energy source. At the onset of cachexia, RER values in tumor-bearing C26 mice fed NF flattened compared to controls on the same diet, indicating that caloric restriction associated with cachexia induces a switch from carbohydrate utilization to exploitation of other energy sources, such as protein or fat stores, that may be responsible for the depletion of muscle and adipose tissue (Figure 6E). Cachectic tumor-bearing C26 mice on KD were unable to adapt their RER and kept using fat reserves, with extremely low RER values around 0.65 and no diurnal fluctuations (Figure 6E). However, Dexamethasone-treated tumor-bearing C26 mice fed KD exhibited a fluctuating metabolism, with fat used as nutrient fuel during light hours but an increase in RER during night time that suggested improved metabolic adaptation and less utilization of fat as energy fuel (Figure 6E). This is in keeping with the observed fat tissue preservation and reduced fatty acid metabolism in these mice (Figure 6D and S5D). Moreover, Dexamethasone treatment increased circulating glucose levels (Figure 6F), hepatic gluconeogenesis and TCA cycle activity (Figure 6G and 6H) in C26 tumor-bearing mice fed KD, therefore improving glucose-homeostasis and whole-body metabolic adaptation and may ultimately be responsible for the observed extended survival of these mice.

Dexamethasone treatment did not change RER levels of C26 mice fed NF, but led to changes in circadian rhythmicity that were almost opposed to littermate controls and untreated C26 mice on NF (Figure 6E). Energy expenditure or heat, referred to the amount of energy (kcal) used to maintain essential body functions per hour, was significantly diminished in cachectic mice compared to their counterparts on the same diet. Dexamethasone treatment markedly raised the energy expenditure of tumor-bearing C26 mice on KD, indicating a beneficial effect of Dexamethasone on the metabolic homeostasis of tumor-bearing mice fed KD, as opposed to C26 tumor-bearing mice fed NF that showed unchanged energy expenditure upon treatment with Dexamethasone (Figure 61). Dexamethasone treatment also significantly enhanced food and water intake in tumor-bearing C26 mice fed either diet to levels higher than their untreated tumor-bearers (Figures 6J and S5F). This anorexia-blocking effect is independent of the metabolic benefits of Dexamethasone, which are exclusive to KD-fed mice. Tumor-bearing mice minimized their total movement in all axis as they developed cachexia, but Dexamethasone treatment increased the total activity of tumor-bearing C26 mice on KD compared to their untreated counterparts. Dexamethasone did not have an effect on the movement of C26 mice fed NF (Figure S5G).

In combination, these results are consistent with a systemic role of glucocorticoids in counteracting cachexia-induced metabolic stress by coordinating adaptive metabolic, feeding and behavioral responses in order to promote survival. Furthermore, the well-known immunosuppressive side effect of glucocorticoid drugs becomes detrimental in the context of a carbohydrate-based diet because it prompts immune escape (Flint et al. 2016) and consequently rapid tumor growth, yet this is overcome in a lipid-enriched, low-carbohydrate diet that induces ferroptotic death of cancer cells. Altogether, this supports a potential synergistical benefit of combining cancer-targeted nutritional interventions with systemic approaches that ameliorate cancer cachexia.

## DISCUSSION

In this study, we find that KD administration to murine models bearing interleukin-6 associated tumors that reduce hepatic ketogenesis (Flint et al. 2016), uncouples tumor progression from overall survival. We trace this to a biochemical sequence where fat-enriched diets enhance the production of lipid-derived reactive molecules that saturate the GSH pathway and deplete NADPH stores. This induces elevations of the stress-induced suppressant of appetite GDF-15. In addition, insufficiency of NADPH cofactor leads to a biochemically-induced relative hypocorticosteronemia and, consequently, metabolic maladaptation in response to stress and earlier onset of cancer cachexia. Dexamethasone administration improves food intake, systemic metabolism, and ultimately, extends survival of tumor-bearing mice fed KD. The relative adrenal insufficiency may only be subclinical in humans, because our preclinical study shows lack of an appropriate upregulation of cortisol in the context of metabolic stress, not a complete absence of the hormone. Thus, even if not presenting apparent symptoms, HPA axis activity and adrenal gland responsiveness should be assessed in patients with a lipid-rich nutritional intake, since this will have effects in systemic metabolism and therapeutic outcome.

These findings caution against a universal utilization of delayed tumor growth as a predictor of prolonged overall survival in cancer models and in the clinic. They highlight that the homeostatic control of energy balance is a highly evolved biological process that involves the coordinated regulation of food intake and energy expenditure. Disruption in metabolic homeostasis causes poor prognosis and early death of patients with cancer (Hursting and Berger 2010), suggesting that therapeutic support of host metabolic adaptation may extend lifespan. Indeed, we find that rescue of this systemic metabolic adaptation (KD plus Dexamethasone) suppresses cancer cachexia and extends survival without altering tumor burden, indicating that it is the systemic metabolic imbalance that is most lethal. At the same time, targeting metabolic dependency of cancer cells shows therapeutic promise, in that it can stall or delay tumor growth. However, our results show that careful consideration of this paradigm is indicated, if a chosen nutritional intervention challenges both the organism and the cancer cell metabolism. This is specifically the case for ketogenic diet interventions, which are currently tested in clinical trials. Here, the anti-cancer effect may be offset by the inability of the organism to utilize the fatty acid nutrients, because reprogramming of systemic metabolism, muscle and fat loss, and reduced food intake are hallmarks of organisms with cancer progression and cancer cachexia (Janowitz 2018).

Glucocorticoids, and specifically cortisol regulate metabolism in conditions of stress. Cortisol increases the availability of blood glucose to the brain and stimulates fat and carbohydrate metabolism. Dexamethasone is a corticosteroid commonly used as a supportive care co-medication for patients with cancer undergoing standard care in order to lower the immune response and reduce inflammation. It is also used as first-line single agent or in combination to prevent or treat cancer-related conditions such as anemia, cerebral oedema, hypersensitivity, hypercalcemia and thrombocytopenia. Side effects of Dexamethasone treatment include weight gain, increased glucose levels and fat accumulation. However, our study shows that what would normally be considered in the clinic as metabolic aftereffects of Dexamethasone may become beneficial for pre-cachectic organisms on a fat-rich diet that exhibit metabolic imbalance and inadequate response to preserve glucose homeostasis (e.g., HPA axis unresponsiveness).

Moreover, some preclinical studies and clinical trials suggest that corticosteroid-induced immunosuppression might dampen the activity of cancer chemo-immunotherapy and increase risk of cancer recurrence (Arbour et al. 2018; Eggermont et al. 2020; Fucà et al. 2019), but this is contradicted by others (Massucci et al. 2020; Menzies et al. 2017). This ongoing debate on the potential impact of corticosteroids on the anti-tumor immune response is illustrated in our preclinical work, as we observe that Dexamethasone administration reduces PFS in mice fed NF while tumor growth is unaffected by Dexamethasone treatment in mice fed KD. Thus, the patients’ dietary intake and nutritional state may be cofounding factors in clinical trials that investigate the immunosuppressive impact of corticosteroids co-treatment in cancer. Nutritional interventions combined with the minimum effective dose should be preferred in the context of corticosteroid treatment.

Dietary interventions can be used to enhance anticancer therapy and improve clinical outcomes (de Groot et al. 2020; Hopkins et al. 2018; Maddocks et al. 2017). Cancer cells exhibit a high rate of glycolysis in the presence of ample oxygen, a process termed aerobic glycolysis (Warburg effect) (Warburg 1925), and this glucose-dependency of tumors can be exploited by specific diet regimens depleted of the nutrients that tumors use as energy source. Diets can also have an effect on the immune system and influence the anti-tumor response. KDs have been previously studied in patients with cancer, and they have been shown to be safe, feasible, and even to have anticancer effects. These low-carbohydrate diets reduce circulating glucose levels and therefore, successfully starve tumors. In this study, we show evidence that supports that KD may be slowing down tumor growth not only via glucose deprivation of the tumor but also through an ongoing accumulation of non-detoxified highly-reactive lipid peroxidation products that causes ferroptotic cell death within the tumor. KD’s aminoacid composition may also be contributing to the observed ferroptosis and metabolic phenotype, as the 9-fold decrease in cystine in the nutritional profile of KD compared to NF can be linked to an induction of ferroptosis independently of fatty acids (Badgley et al. 2020). Changes in metabolism and energy expenditure in KD-fed mice could be partially attributed to the 3-fold decrease in methionine intake (Pissios et al. 2013) (Extended Data Table 1).

We note that the effect of KD on the host is likely context-dependent: it is detrimental for both the tumor and the host that has been metabolically reprogrammed and bears an established tumor, uncoupling tumor size from overall survival; but KD has no impact on non­tumor-bearing hosts that are able to adapt their metabolism to the nutritional intake. Even if they grow a tumor later on, the metabolic reprogramming induced by the tumor is delayed due to its decelerated growth. This explains differential results in the literature (Lien et al. 2021) and evidences the importance of dosing and timing in the clinic.

Without independent expansion and validation of our results, it is perhaps too early to suggest that KD and glucocorticoid co-administration may be a therapeutic strategy for patients with interleukin-6 elevating cancers. Nevertheless, the comparatively reduced tumor progression and prolonged overall survival in mice on KD and Dexamethasone compared to mice on NF with or without Dexamethasone is an encouraging finding. A limitation to this concept and to our study is a lack of the exact understanding of how glucocorticoids reprogram and rescue metabolism in the context of a reprogrammed organism challenged with high lipid diet. While this may limit the ability for longitudinal molecular response monitoring of patients on the combination intervention, weight trajectories and glucose levels may be suitable and readily obtainable biomarkers for clinical studies.

## CONCLUSION

Our study highlights that the effect of systemic interventions cannot necessarily be extrapolated from the effect on the tumor alone, but that they have to be investigated for anti­cancer and host effects. In model systems with established tumors that elevate interleukin 6, the opposing effect of a KD nutrition on delayed tumor growth and induction of cachexia, lead to a dominant negative effect on overall survival. These findings may be relevant to clinical research efforts that investigate the potential benefit of KD for patients with cancer.

## LIMITATIONS

Our results have been obtained using model systems of colorectal and pancreatic cancer that are known to recapitulate clinical disease progression from early cancer to cancer cachexia, but clinical validation of our work is needed. We acknowledge that not all cancers lead to IL-6 elevations and we cannot comment on the transferability of our findings to cancers that are not associated with raised IL-6. The time-course of disease progression and metabolic reprogramming in patients with cachexia inducing tumors has not been resolved with high resolution. We here focus on mice with fully established tumors that are challenged with KD when early metabolic reprogramming has occurred. Future studies have to guide how preclinical work like this is best translated to clinical cancer progression and how this alignment could guide stratified enrolment of patients into interventional trials. Our work also provides no definitive answer on dosing for glucocorticoid replacement and, here too, detailed clinical studies are required to define the best therapeutic dose range.

## Supporting information

Key Resources Table

## ACKNOWLEDGMENTS

Miriam Ferrer is supported by the “la Caixa” Foundation (ID 100010434) in the framework of the “La Caixa” Fellowship Program under agreement LCF/BQ/AA18/11680037, and by the MRC Cancer Unit with a Doctoral Training Award. Tobias Janowitz acknowledges funding from Cancer Grand Challenges (NIH: 1OT2CA278690-01; CRUK: CGCATF-2021/100019), Cancer Research UK (C42738/A24868), The Mark Foundation for Cancer Research (33300111), Cold Spring Harbor Laboratory (CSHL), and developmental funds from CSHL Cancer Center Support Grant 5P30CA045508. The CRUK CI (Li Ka Shing Centre) where some of this work was performed was generously funded by CK Hutchison Holdings Limited, the University of Cambridge, CRUK, The Atlantic Philanthropies and others. This work was supported by Medical Research Council (MRC) Programme grants MC_UU_12022/l and MC_UU_12022/8 to Ashok R. Venkitaraman.

## AUTHOR CONTRIBUTIONS

Conceptualization, M.F. and T.J..; Methodology, M.F., A.R.V., T.J.; Investigation, M.F., N.M., E.E.D., S.O.K., M.Z., J.H, R.R., T.R.F., C.M.C., and M.L.; Writing - Original Draft, M.F.; Writing - Review & Editing, all authors.; Project Administration, M.F; Funding Acquisition, M.F., A.R.V. and T.J.; Resources, A.R.V. and T.J.; Supervision, T.J.

## STAR METHODS

### Laboratory animals

Two different mouse models that predispose to cachexia were used. A transplanted C26 model of colorectal cancer and the genetically engineered autochthonous KPC model of pancreatic cancer. The C26 model is performed on wild-type BALB/c mice that are inoculated subcutaneously with a syngeneic tumor. In the KPC system, an activating point mutation (G12D) in Kras and a dominant negative mutation in Trp53 (R172H) are conditionally activated in the pancreas by means of the Cre-Lox technology. Both pre-clinical models have been shown to develop tumors that secrete IL-6 and therefore the host is unable to produce ketones during caloric deficiency associated with cachexia, causing a rise in glucocorticoid levels as a consequence. KPC and BALB/c mice were obtained from Charles River Laboratories. They were kept in pathogen-free conditions on a 24 hour 12:12 light-dark cycle and allowed to acclimatize for 7 days. All animal experiments and animal care at the MRC CU and CRUK CI were performed in accordance with national and institutional guidelines and approved by the UK Home Office, the animal ethics committee of the University of Cambridge. All animal experiments at CSHL were approved by the Institutional Animal Care and Use Committee (IACUC) and were conducted in accordance with the National Institutes of Health Guide for the Care and Use of Laboratory Animals. Body weights, food intake and clinical signs were monitored on a daily basis. Handling was kept to a minimum. Mice were described as cachectic when they reached weight loss of more than 5% from their peak weight and were sacrificed when tumor size exceeded 2 cm length, when weight loss exceeded 15% from peak weight, or when showing clinical signs of discomfort indicative of cachectic endpoint as stated by the Animal Cachexia Score (ACASCO): piloerection, diarrhea or constipation, hunched posture, tremors, and closed eyes (Betancourt et al. 2019). Death was confirmed by cervical dislocation.

#### Experimental enrolment

Weight-stable, tumor-bearing male KPC mice with tumors of 3-5 mm size and no evidence of obstructive common bowel duct, and their respective weight- and age-matched control counterparts (PC mice) were enrolled in the experiments.

Weight-stable wild-type 9-weeks old male BALB/c mice were inoculated with 2xl0^6^ viable C26 colorectal cancer cells subcutaneously (s.c.) and enrolled on study together with their respective controls.

At day of enrolment (day 14 post-injection of C26 cells), mice were stratified in terms of tumor size, body weight and age, singly-housed and randomly allocated into two experimental matched groups fed with different diets: mice were fed with either standard diet (#5053 PicoLab® Rodent Diet 20; LabDiet) or ketogenic diet (KD) (#F3666; Bio-Serv). Ketogenic diet was given in a Petri dish container that was replaced daily due to potential oxidation of the diet.

#### Endpoint

Overall survival (OS) or Progression Free Survival (PFS) were the final endpoint of the studies.

#### Overall Survival

Mice were considered to have reached the OS endpoint when their body weight loss exceeded 15% from their peak weight.

#### Progression Free Survival

Mice were considered to have reached the PFS endpoint when the volume of their tumors exceeded 2000 mm^3^, as measured by handheld calipers.

#### Dexamethasone treatment

Dexamethasone 21-phosphate disodium salt (#D1159; Sigma-Aldrich) was dissolved in dl·l_2_O and administered intraperitoneally (i.p.) at 1mg/kg daily.

#### N-acetyl cysteine (NAC) treatment

N-Acetyl-L-cysteine (#A9165; Sigma-Aldrich) was dissolved in 0.9% NaCl sterile saline solution (#Z1377; Thermo Fisher) and administered intraperitoneally (i.p.) at 150mg/kg daily.

#### RSL3 treatment

RSL3 ((1S,3R)-RSL3) (#HY-100218A; MedChemExpress) was dissolved in 10% Dimethyl Sulfoxide (DMSO) (#12611S; Cell Signaling), 40% Polyethylene glycol 300 (PEG300) (#S6704; Selleck Chemicals), 5% Tween-80 and 45% NaCl 0.9% sterile saline solution (#Z1377; Thermo Fisher) and administered intraperitoneally (i.p.) at 5mg/kg daily.

#### ACTH stimulation test (synacthen test)

ACTH (#HOR-279, ProSpec) was reconstituted in dl·l_2_0 and injected intraperitoneally at a dose of 1 ug/g body weight. Tail blood was collected at 0-, 15-, 30-, and 60-min intervals for determination of plasma corticosterone levels. Each group consisted of five animals.

### Metabolic cages

The Comprehensive Lab Animal Monitoring System (CLAMS) from Columbus Instruments was used to monitor and quantify multiple metabolic parameters such as activity, weight (g), drinking (mL), food intake (g), sleep, body core temperature and open circuit calorimetry in animal cages that allows precise control over the light/dark cycle. Data from C26- injected or WT BALB/c control mice on standard or ketogenic diet treated with Dexamethasone or vehicle (saline) was collected real-time during an acclimation period (72 hours), a baseline period (72 hours) and during all the experimental timeline (approximately 33 days) through the Oxymax collection software during 10-30 second intervals. Data was exported and analyzed in RStudio.

### Tumor size

PDAC tumors in KPC mice were detected via palpation and high-resolution ultrasound imaging (Vevo 2100; VisualSonics), and confirmed at necropsy. Tumor growth was monitored by ultrasound scans assessed at multiple angles. Mice were carefully observed for any macroscopic metastases. Maximum cross-sectional area (CSA) and maximum diameter of the tumors were determined for each timepoint. Tumor development in BALB/c mice was spotted via palpation and monitored daily by caliper measurements. Volume of the tumor was calculated as follows: volume (mm^3^) = [long axis (mm) x short axis (mm)]^2^/ 2.

### Blood and plasma measurements

Tail bleeds and terminal cardiac bleeds were taken. Tail vein bleeds were performed using a scalpel via tail venesection without restraint, and terminal bleeds were obtained through exsanguination via cardiac puncture under isoflurane anesthesia. Tail bleeds were immediately analyzed for glucose and ketone concentration measurements using glucose/ketones stripes and gluco-/keto-meters (Freestyle Optium Neo; Abbott laboratories).

Plasma samples were collected from tail or terminal cardiac bleeds using heparin-coated hematocrit capillary tubes to avoid coagulation and were processed as follows: centrifuge spin at 14,000 rpm for 5 min at 4°C, snap frozen in liquid nitrogen and stored at -80°C.

Corticosterone was quantified from plasma using the International Corticosterone (Human, Rat, Mouse) ELISA (#RE52211; IBL). The sample incubation step from the IBL assay protocol was 3 hours at room temperature (RT) so as to reach displacement equilibrium as determined by preliminary data. IL-6 levels were measured from plasma using the mouse IL-6 Quantikine ELISA Kit (#M6000B; R&D Systems).

4-hyroxynonenal (4-HNE) measurements in the plasma were quantified using the Universal 4-Hydroxynonenal ELISA Kit (Colorimetric) (#NBP2-66364; Novus Biologicals).

Pregnenolone was measured in the plasma using the Pregnenolone ELISA Kit (Colorimetric) (#NBP2-68102; Novus Biologicals).

Levels of Adrenocorticotropin hormone (ACTH) in the plasma were measured with the Mouse/Rat ACTH ELISA Kit (#ab263880; Abcam).

### Tissue collection

Liver, tumor, spleen, adrenal glands, quadricep muscle, lungs, and gonadal and subcutaneous fat samples were collected and weighed during necropsy dissection. Subsequently, tumor, liver and spleen samples were cut into 3 equal parts, which were either snap frozen in liquid nitrogen, cryo-embedded in OCT, or fixed in 4% neutral buffered formaldehyde for 24 hours at room temperature (RT) before either being transferred to 70% ethanol and later paraffin-embedded (FFPE) for immunohistochemistry processing. All the other organs and tissue samples were immediately snap frozen and stored at -80°C.

### Tissue lysis

Snap frozen tissues stored at -80°C were transferred to dishes on wet ice and cut into pieces with a scalpel. Each piece was weighed and placed into 2mL round-bottom Eppendorf tubes pre-loaded with Stainless Steel beads (#69989; Qiagen) on wet ice. Homogenizer tubes were then filled up with lysis buffer (#AA-LYS-16ml; RayBiotech) and supplemented with

Protease Inhibitor Cocktail (#AA-PI; Raybiotech) and Phosphatase Inhibitor Cocktail Set I (#AA- PH1-1; RayBiotech). Samples were homogenized in Tissue Lyser II (#85300; Qiagen) for 5 minutes and then lysates were centrifuged at 4°C for 20 minutes at maximum speed. The supernatant was harvested and kept on ice if testing fresh or sored at -8O°C.

The Bicinchoninic Acid (BCA) Method was used to determine protein concentration in lysates.

Ferrous (Fe^2+^) and Ferric (Fe^3+^) iron levels in tissue lysates were measured using the Colorimetric Iron Assay Kit (#ab83366; Abeam).

Quantification of 4-HNE-protein adducts in lysates was performed using the Lipid Peroxidation (4-HNE) Assay Kit (#ab238538; Abeam).

Detection of NADPH levels in the adrenal glands was performed using the NADP/NADPH-Glo™ Bioluminescent Assay (#G9081; Promega).

### Oil-Red-O staining

Fresh frozen tissue sections of 5-10 µm thickness were mounted on slides, air dried for 30-60 minutes at RT and fixed in ice cold 10% neutral-buffered formalin (#HT501128-4L; Sigma- Aldrich) for 5-10 minutes. After rinsing in 3 changes of distilled water and air drying for another 30-60 minutes, slides were placed in absolute Propylene Glycol (#P4347; Sigma-Aldrich) for 2-5 minutes, then stained in pre-warmed Oil Red O solution (#O0625-25G; Sigma-Aldrich) for 8-10 minutes in a 60°C oven, differentiated in 85% Propylene Glycol solution for 2-5 minutes and rinsed in 2 changes of distilled water. Slides were then counterstained with Mayer’s hematoxylin (#ab245880; Abcam) for 30 seconds, washed thoroughly with distilled water and mounted with aqueous mounting medium.

### Immunohistochemistry

Tissues were fixed in 4% paraformaldehyde (#50-980-495; Thermo Fisher) for 24 h and then embedded in a paraffin wax block. Sectioning with a cryostat, deparaffinization, antigen retrieval and immunohistochemistry for Kİ67 (#14-5698-82; Thermo Fisher) was performed by the CRUK Cl Histopathology Core using a Leica Bond III autostainer. The slides were scanned on a Leica Aperio AT2 system and subsequently analyzed in a blinded manner. H&E staining was performed by the Histology Facility at Cold Spring Harbor Laboratory.

### Metabolomics

Global metabolic profiling of liver and tumor samples was performed by UPLC-MS/MS at Metabolon, Inc. facilities (UK project #CRUK-01-19VW; USA project #CSHL-01-22VW+; results from both datasets were merged). Samples were prepared using the automated MicroLab STAR® system from Hamilton Company. Several recovery standards were added prior to the first step in the extraction process for QC purposes. To remove protein, dissociate small molecules bound to protein or trapped in the precipitated protein matrix, and to recover chemically diverse metabolites, proteins were precipitated with methanol under vigorous shaking for 2 min (Glen Mills GenoGrinder 2000) followed by centrifugation. The resulting extract was divided into five fractions: two for analysis by two separate reverse phases (RP)/UPLC-MS/MS methods with positive ion mode electrospray ionization (ESI), one for analysis by RP/UPLC-MS/MS with negative ion mode ESI, one for analysis by HILIC/UPLC-MS/MS with negative ion mode ESI, and one sample was reserved for backup. Samples were placed briefly on a TurboVap® (Zymark) to remove the organic solvent. The sample extracts were stored overnight under nitrogen before preparation for analysis.

Raw data were extracted, peak-identified and QC processed using Metabolon’s hardware and software. These systems are built on a web-service platform utilizing Microsoft’s .NET technologies, which run on high-performance application servers and fiber-channel storage arrays in clusters to provide active failover and load-balancing. Compounds were identified by comparison to library entries of purified standards or recurrent unknown entities. Metabolon maintains a library based on authenticated standards that contains the retention time/index (RI), mass to charge ratio (m/z), and chromatographic data (including MS/MS spectral data) on all molecules present in the library. Furthermore, biochemical identifications are based on three criteria: retention index within a narrow RI window of the proposed identification, accurate mass match to the library +/- 10 ppm, and the MS/MS forward and reverse scores between the experimental data and authentic standards.

A total of 685 and 669 named metabolites were retained for liver and tumor datasets, respectively. Following log transformation and imputation of missing values, if any, with the minimum observed value for each compound, Welch’s two-sample t-test was used to identify biochemicals that differed significantly between experimental groups. An estimate of the false discovery rate (q-value) was calculated to take into account the multiple comparisons that normally occur in metabolomic-based studies.

### Cell lines

#### C26 murine colorectal cancer cell line

C26 cells were cultured in complete growth medium consisting of RPMI-1640 medium with Glutamine (#11-875-093; Thermo Fisher) containing 10% of Heat-Inactivated Fetal Bovine Serum (FBS) (#10-438-026; Thermo Fisher) and lx Penicillin- Streptomycin solution (#15-140-122; Thermo Fisher) under sterile conditions, lx Trypsin-EDTA (#15400054; Thermo Fisher) was used for cell dissociation. Cells were resuspended in FBS-free RPMI and viable cells were counted using a Vi-Cell counter prior to subcutaneous injection of 2xl0^6^ viable cells diluted in lOOµL RPMI into the right flank of each BALB/c mouse.

#### H295R human adrenocortical cell line

H295R cells (#CRL-2128; ATCC) were cultured in complete growth medium consisting of DMEM:F12 medium (#30-2006; ATCC) with 0.00625 mg/mL insulin; 0.00625 mg/mL transferrin; 6.25 ng/mL selenium; 1.25 mg/mL bovine serum albumin; 0.00535 mg/mL linoleic acid (ITS+ Premix)(#354352; Corning) and adjusted to a final concentration of 2.5% Nu-Serum I (#355100; Corning). Cells were grown in 75 cm^2^ culture flasks and subcultured 1:3 every 3 days, lx Trypsin-EDTA solution was used for cell dissociation prior to seeding cells in 96-well plates for experimental viability or cortisol release tests.

#### H295R Cortisol synthesis assay

H295R cells were seeded in 96-well plates with 12,000 cells in 200 µL of complete growth medium for each well and allowed to settle for 48 hours. The medium was then changed and 200 µL medium containing specific LPPs was added. LPPs used: 4-hydroxynonenal (4-HNE) (#32100; Cayman Chemical), 4-hydroxyhexenal (4-HHE) (#32060; Cayman Chemical) and malondialdehyde (MDA) (#63287-1G-F; Sigma-Aldrich). After 48 or 72 hours of incubation in the presence of the compound, the medium was carefully collected, transferred to the test tubes and immediately stored at -20°C. Cells in each well were detached and counted by Trypan Blue staining (#15250061; Thermo Fisher). Samples were diluted 1:5 and cortisol levels were measured using a Cortisol Competitive Human ELISA Kit (#EIAHCOR; Thermo Fisher) according to the manufacturer’s protocol.

### Sulforhodamine B (SRB) colorimetric assay

A total of 12,000 cells per well were seeded in a 96-wells plate. Two days after seeding, media was replaced and cells were treated with increasing concentrations of a specific LPP for 72h. Then, cells were fixed with 1% trichloroacetic acid (#T9159-100G; Sigma-Aldrich) at 4I3°C for 30 min, washed 3 times with distilled water and stained with 0.057% SRB (#S1402-5G; Sigma-Aldrich) in 1% acetic acid (#A6283, Sigma-Aldrich) solution at RT for 30 min. Following staining, cells were washed 3 times with 1% acetic acid and air-dried overnight. The protein bound dye was dissolved in lOĒmM Tris base solution and the absorbance was measured at 565lānm using a microplate reader (SpectraMax İ3x).

### Single cell preparation

Cell suspensions were prepared from tissues by mechanical dissociation, followed by digestion in 5 mL of RPMI-1640 containing collagenase I (500 U/mL) (#SCR103; Sigma-Aldrich) and DNase I (0.2 mg/mL) (#04716728001; Sigma-Aldrich) for 45 min at 37°C on a shaker (220 rpm), followed by filtration through a 70-µm strainer and 25% Percoll (#GE17-0891-01; Sigma-Aldrich) gradient enrichment of leukocytes, and red blood cell (RBC) lysis. Tumor cells were recovered without Percoll enrichment. Blood cells were lysed in 5 mL of RBC lysis buffer (#A1049201; Thermo Fisher) three times for 5 min, and spleens were strained through a 70-µm filter in RPMI-1640 before lysing erythrocytes with RBC lysis buffer for 5 min. Single cells were restimulated and stained for surface and intracellular markers (see flow cytometry below).

### Flow cytometry

Cell sorting was performed using a FACSAria™ Cell Sorter (BD Biosciences) at CSHL Flow Cytometry Facility. FlowJo X (Tree Star) software was used for experimental analysis.

The following antibodies were used: Alexa Fluor 700 anti-mouse CD45 (#103127; BioLegend), FITC anti-mouse CD45 (#11-0451; Thermo Fisher), APC/Cy7 anti-mouse CD3ε (#100329; BioLegend), PerCP/Cyanine 5.5 anti-mouse CD4 (#100433; BioLegend), Brilliant Violet 510TM anti-mouse CD8a (#100751; BioLegend), Brilliant Violet 605TM anti-mouse/human CDllb (#101257; BioLegend), Alexa Fluor 700 anti-mouse Ly-6G/Ly-6C (Gr-1) (#108421; BioLegend), FITC anti-mouse CD69 (#104505; BioLegend), PE/Cy7 anti-mouse CD152 (#106313; BioLegend); Brilliant Violet 421TM anti-mouse CD274 (#124315; BioLegend), PE/Dazzle 594 anti-mouse CD279 (#109115; BioLegnd), and FITC anti-mouse F4/80 (#123107; BioLegend).

### Western blotting

Cells were lysed in RIPA buffer (50mM Tris HCl, pH 7.4, 150mM NaCl, 0.5% deoxycholate, 0.1% sodium dodecyl sulphate, 1% NP-40) (#89901; Thermo Fisher) containing protease inhibitors (#78442; Thermo Fisher) and 1 mM dithiothreitol (DTT) (#A39255; Thermo Fisher). Whole cell extracts were separated by electrophoresis, transferred onto nitrocellulose membranes (#88025; Thermo Fisher) and blocked in 5% non-fat dry milk (#1706404; Bio-Rad) dissolved in 0.1% Tween/TBS. Membranes were incubated with primary antibodies: BAX Rabbit mAb (#50599-2-Ig; Proteintech, 1:500), β-Actin Rabbit mAb (#4967; Cell Signaling Technology, 1:5000) and Caspase-3 (D3R6Y) Rabbit mAb (#14220; Cell Signaling, 1:1000), overnight at 4°C followed by washing in 0.1% Tween/TBS. Membranes were incubated with Goat Anti-Rabbit IgG H&L (HRP) secondary antibodies (#ab205718; Abcam, 1:5000) at 25°C for 1h and washed thrice prior to signal detection. Membranes were developed by exposure in a dark room through chemiluminescence using ECL reagent (#32106; Thermo Fisher).

### qRT-PCR

mRNA was extracted from frozen tissues using QIAzol Lysis Reagent (#79306; Qiagen) and the Tissue Lyser II (#85300; Qiagen), following the manufacture’s protocol for the RNeasy Lipid Tissue Mini Kit (#74804, Qiagen) in an automated manner with the QIAcube Connect (#9002864; Qiagen). Concentration and purity of aqueous RNA was assessed using a NanoDrop™ Spectrophotometer (#ND-ONE-W; Thermo Fisher). mRNA templates from muscle and liver samples were diluted to 2ng/µl and mRNA was analysed by quantitative Real-Time PCR using the TaqMan™ RNA-to-CT™ 1-Step Kit (#4392653; Thermo Fisher). mRNA levels were normalized to either Rnlδs (liver) or Tbp (quadriceps) using the ddCt method. The following TaqMan primers were used: Mm01277044_m1 (Tbp); Mm03928990_gl (Rnlδs); Mm00440939_m1 (Ppara); Mm01323360_gl (Acadm); Mm00550050_m1 (Hmgcs2); Mm00499523_ml (Fbxo32); and Mm01185221_m1 (Trim63).

### RNA-sequencing

RNA extracted from frozen tissues via QIAzol Lysis Reagent (#79306; Qiagen) was run through RNeasy spin columns following the RNeasy Lipid Tissue Mini Kit in an automated manner with the QIAcube Connect (#9002864; Qiagen). Integrity was confirmed using RIN values with a cut-off of 8. Libraries were prepared by the Next Gen Sequencing Core at CSHL using the lllumina TruSeq mRNA Stranded Sample prep kit (96 index High Throughput) and normalized using Kapa Biosystem’s Library Quantification Kit. NextSeq High Output Paired-End 150bp was run for sequencing.

For the analysis, reads were aligned to the mouse genome version GRCm38.74 and read counts were obtained using “biomaRt” R package. “org.Mm.eg.db” R package was used for genome wide annotation. Read counts were normalized and tested for differential gene expression using the Bioconductor package “edgeR”. Multiple testing correction was applied using the Benjamini-Hochberg procedure (FDR <0.05). “fgsea” and “dplyr” packages were used for GSEA in R. GSEA was performed by ranking all genes tested in RNA-Seq using -loglO (p- values) derived from differential expression analyses and testing against MSigDB Hallmark gene sets and Canonical pathways KEGG gene sets. Results were curated using a p-adj <0.05 threshold.

### Statistical analysis

Data were expressed as the mean ± SEM unless otherwise stated and statistical significance was analyzed using GraphPad Prism 7.03 software. For survival analysis, data were shown as Kaplan Meier curves and the log-rank (Mantel-Cox) test was used to assess survival differences. When comparing more than 2 groups at the same time, one-way ANOVA with Tukey’s correction for post-hoc testing was used. For statistical comparison of quantitative data at different times, unpaired two-tailed Student’s t-tests were performed at each timepoint with the Holm-Sidak method correction for multiple comparisons. To analyze the main independent effect of diet and cancer and the interaction of both factors, two-way ANOVA tests were used, as well as Welch’s t-test to compare two-samples with different variance.

For global metabolic profiling of liver and tumor, a Principal Component Analysis (PCA) was performed. PCA is an unsupervised statistical method that reduces the dimension of the data by using an orthogonal transformation to convert a set of observations of possibly correlated variables into a set of variables called principal components. Each principal component is a linear combination of every metabolite and the principal components are uncorrelated. The number of principal components is equal to the number of observations This method permits visualization of how individual samples in a dataset differ from each other. The first principal component is computed by determining the coefficients of the metabolites that maximizes the variance of the linear combination. The second component finds the coefficients that maximize the variance with the condition that the second component is orthogonal to the first. The third component is orthogonal to the first two components and so on. The total variance is defined as the sum of the variances of the predicted values of each component (the variance is the square of the standard deviation), and for each component, the proportion of the total variance is computed. Samples with similar biochemical profiles cluster together whereas samples with distinct profiles segregate from one another.

Differences in tumor growth were assessed by fitting a mixed effect model with coefficients for the intercept, slope and the difference in the slope between diets, and a random component for each individual mouse. Significance was assessed by testing whether the coefficient for the difference in the slope was significantly different from zero using a t-test.

## SUPPLEMENTAL INFORMATION TITLES AND LEGENDS

**Figure S1.**
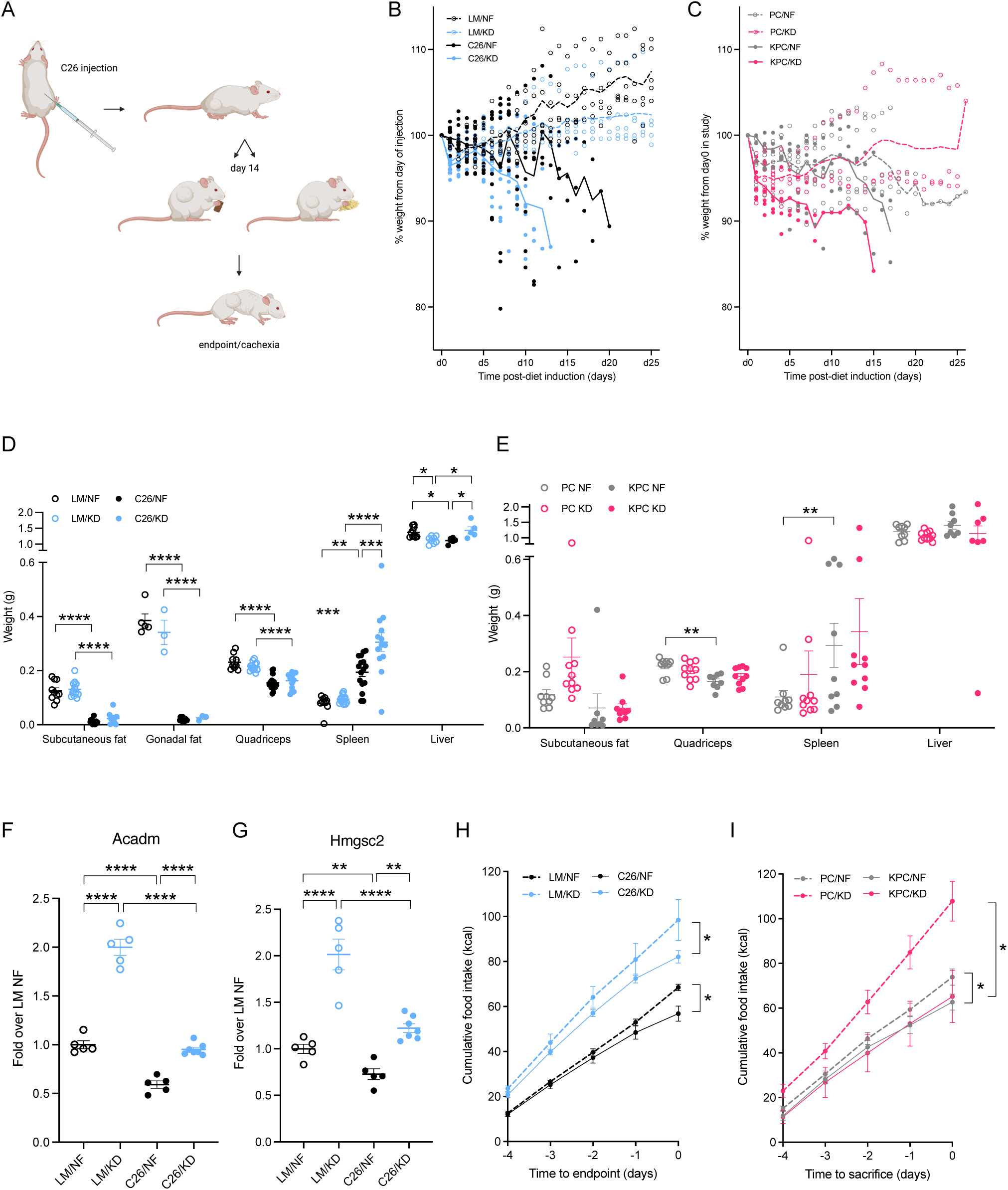
Cachectic phenotype of C26 and KPC murine models. (A) Graphical summary of the experimental protocol. (B-C) Weight trajectories of C26 tumor-bearing mice and littermate controls (C), and KPC tumor-bearing mice and PC controls (C) on KD or NF diets since they were enrolled into the study until they reached cachectic endpoint. (D-E) Organ weights of cachectic C26 tumor-bearing mice and littermates (n=10-15) (D), and cachectic KPC tumor-bearing mice and PC controls (n=9-10) (E) fed either KD or NF diets. (F-G) mRNA expression of the PPARα target genes Acadm (F) and Hmgsc2 (G) in C26 tumor-bearing mice and littermate controls fed KD or NF (n=5-7). (H-I) Cumulative food intake of KD- or NF-fed C26-tumor bearing mice and littermates (n=5 LM, n=12 C26) (H), and KD- or NF-fed KPC tumor-bearing mice and PC controls (n=10) (I) during the last 4 days before endpoint. Data are expressed as the mean ± SEM. One-way ANOVA with Tukey’s correction for post hoc testing was used in (D-G). Two-way ANOVA statistical tests with Tukey’s correction for post hoc comparisons were performed in (H-l). * p-value < 0.05, ** p-value < 0.01, **** p-value < 0.0001.

**Figure S2.**
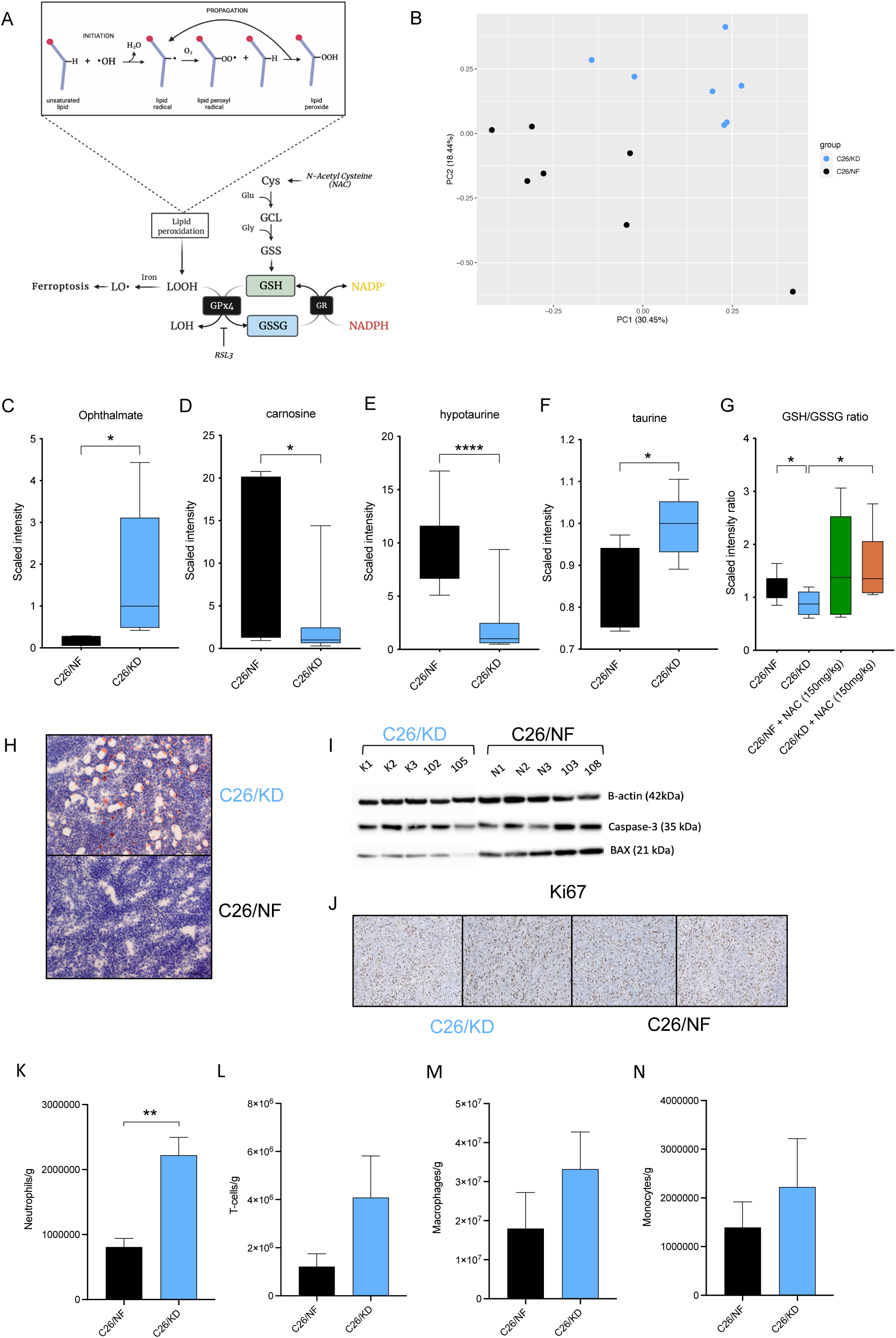
Intratumoral accumulation of lipids and saturation of the GSH system causes ferroptotic cell death. (A) Schematic representation of the GSH pathway for detoxification of LPPs. (B) PCA of tumors from C26 tumor-bearing mice fed NF or KD (n=7). (B) PCA of untargeted metabolomics in the tumors of C26 mice fed KD or NF. (C-F) Quantification of ophthalmate (C), carnosine (D), hypotaurine (E) and taurine (F) metabolites by UPLC-MS/MS in the tumor of C26 mice fed KD or NF (n=7). (G) Quantification by UPLC-MS/MS of GSH/GSSG ratio in the tumor of C26 mice fed KD or NF, untreated or treated with NAC (n=5-7). (H) Oil-Red-O staining of tumors from C26 mice on KD or NF. (I) Western blot of tumor lysates from C26 mice fed KD or NF stained for Caspase-3 and BAX apoptotic markers (n=5). (J) Immunohistochemistry staining of tumors from C26 mice fed KD or NF with the proliferation marker Kİ67. (K-N) Quantification by flow cytometry of neutrophils (K), T-cells (L), macrophages (M) and monocytes (N), in the tumor of C26 mice fed KD or NF (n=3-4). Data are expressed as the mean ± SEM. Statistical analysis in (B) is described in the Methods section/Chapter 4. Statistical differences in (C-F, K-N) were examined using an unpaired two­tailed Student’s t-test with Welch’s correction. One-way ANOVA with Tukey’s correction for post hoc testing was used in (G). * p-value < 0.05, ** p-value < 0.01, **** p-value < 0.0001.

**Figure S3.**
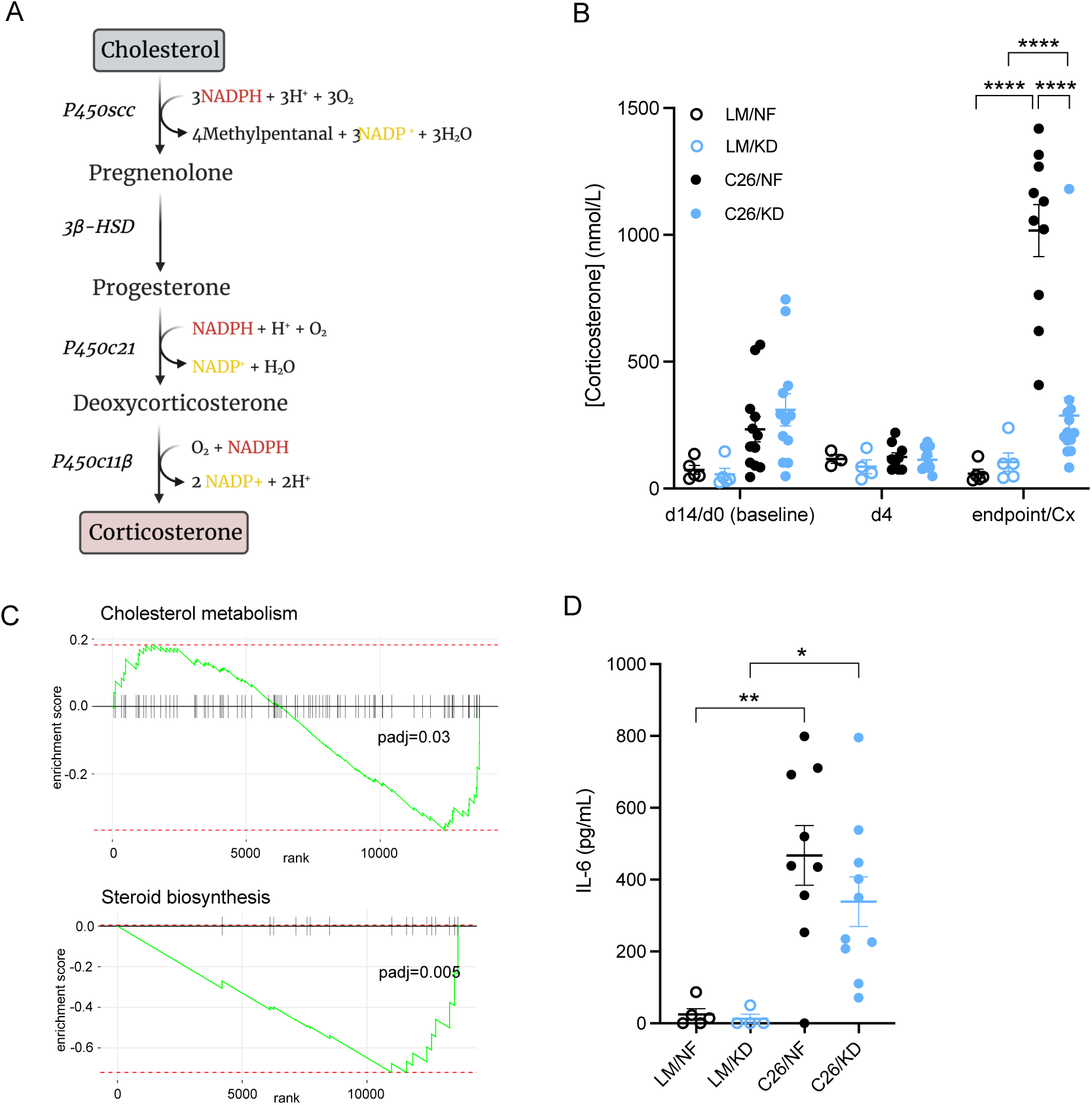
Biochemical deficiency in the corticosterone synthesis pathway in the adrenal cortex of tumor-bearing mice fed ketogenic diet. (A) Murine synthetic pathway of corticosterone in the cortex of the adrenal glands. (B) Corticosterone levels at baseline (prior to diet change), 4 days after the start of the experiment, and at endpoint (cachexia) in C26 tumor­bearing mice and littermate controls fed KD or NF (n=5 LM, n=10-14 C26). (C) GSEA pathway analysis of cholesterol homeostasis and steroid biosynthesis in tumor-bearing KD-fed KPC mice compared to NF-fed KPC (n-5). (D) Plasma concentration of the pro-inflammatory cytokine IL-6 in C26 tumor-bearing mice and control littermates on NF or KD diets at endpoint (n=5 LM, n=9- 10 C26). Data are expressed as the mean ± SEM. One-way ANOVA with Tukey’s correction for post hoc testing was used in (B, D). Statistical analysis in (C) is described in Methods. * p-value < 0.05, ** p-value < 0.01, *** p-value < 0.001, **** p-value < 0.0001.

**Figure S4.**
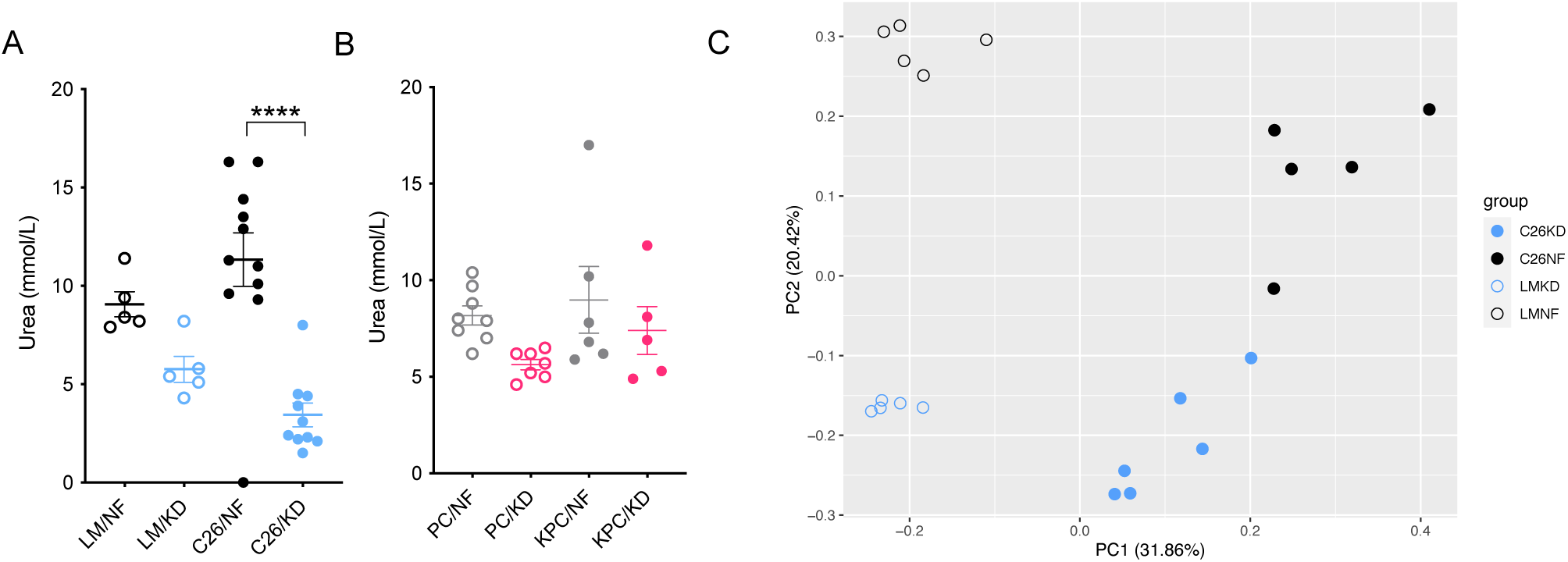
Extended metabolic profiling of cachectic C26 and KPC mice. (A-B) Plasma urea levels in cachectic C26 tumor-bearing mice and littermate controls (n=5 LM, n=10-ll C26) (A), and cachectic KPC tumor-bearing mice and PC controls (n=5-8) (B) fed either KD of NF diets. (C) PCA of hepatic metabolomics in C26 tumor-bearing mice and control littermates fed with KD or NF (n=5-6). Data are expressed as the mean ± SEM. One-way ANOVA with Tukey’s correction for post hoc testing was used in (A-B). Statistical analysis in (C) is described in Methods. * p-value < 0.05, ** p-value < 0.01, **** p-value < 0.0001.

**Figure S5.**
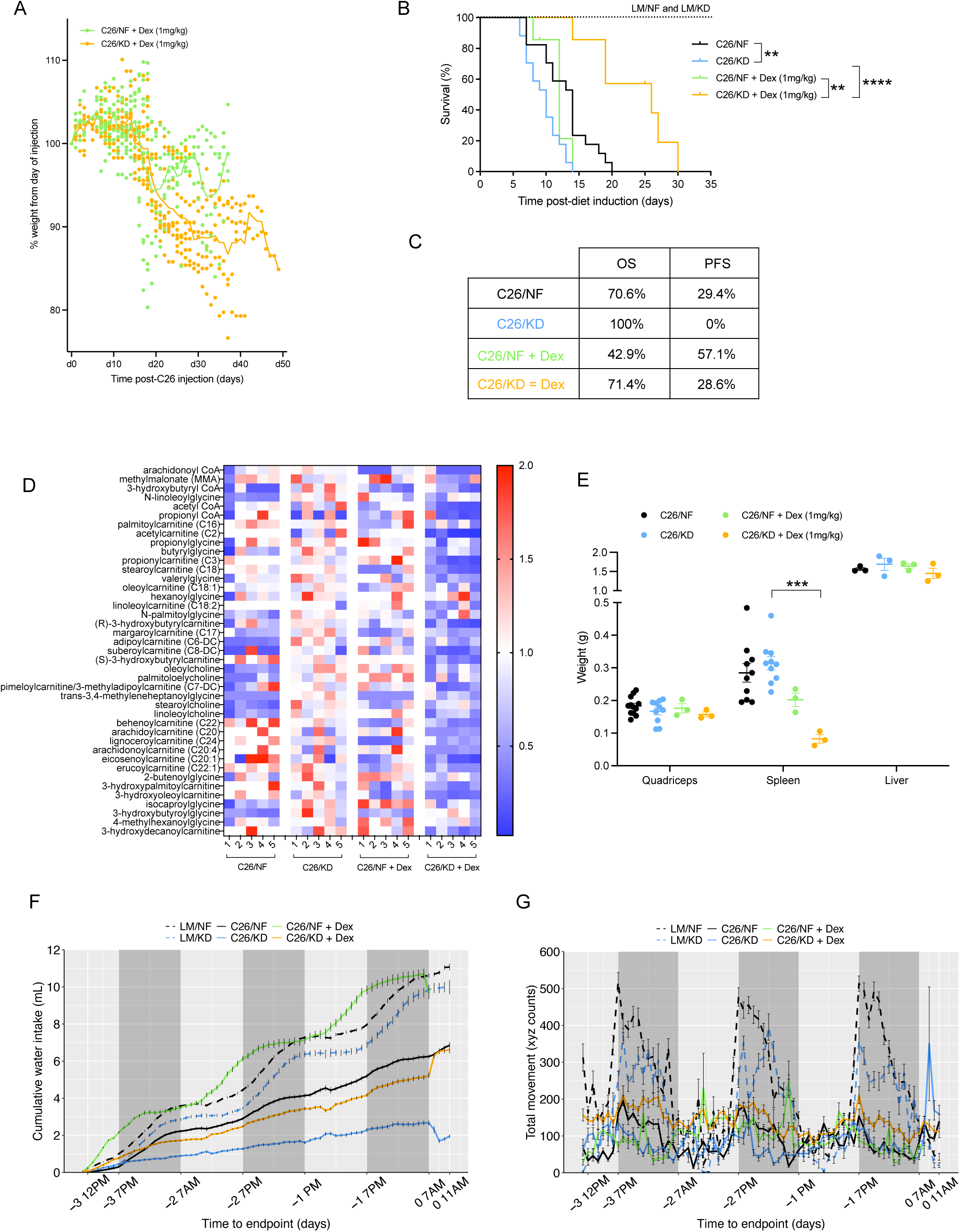
Extended data on the systemic effects of Dexamethasone treatment. (A) Weight trajectories of C26 tumor-bearing mice treated with Dexamethasone and fed with either KD or NF. (B) Survival of littermate controls, and C26 tumor-bearing mice treated or untreated with Dexamethasone, fed with KD or NF (n=7 LM, n=17-18 C26, n=7 C26 + Dex). (C) Percentage of mice in each group that were sacrificed because of cachexia (OS) or tumor size (PFS) endpoints. (D) Quantification by UPLC-MS/MS of metabolites involved in fatty acid metabolism in the liver of C26 tumor-bearing mice on either KD or NF diets, untreated or treated with Dexamethasone (n=5). (E) Organ weights of C26 tumor-bearing mice untreated or treated with Dexamethasone after 4 days of treatment (n=10-12 C26, n=3 C26 + Dex). (F-G) Cumulative water intake (F) and total movement (G) during the last 4 days before endpoint in littermate controls and C26 tumor-bearing mice, untreated or treated with Dexamethasone, fed KD or NF (n=7). Data are expressed as the mean ± SEM. Survival: OS + PFS. Kaplan-Meier curves in (B-C) were statistically analyzed by using the log-rank (Mantel-Cox) test. One-way ANOVA with Tukey’s correction for post hoc testing was used in (E). Analysis in (D, F-G) is described in Methods. * p- value < 0.05, ** p-value < 0.01, *** p-value < 0.001, **** p-value < 0.0001.

**Supplemental Table 1.** Macronutrient composition and caloric profile of standard and ketogenic diets.

|  | Standard diet<br>(PicoLab Rodent Diet 20) | Ketogenic diet<br>(AIN-76A Modified) |
| --- | --- | --- |
| Fat (%) | 10.6 | 75.1 |
| Protein (%) | 20 | 8.6 |
| Carbohydrates (%) | 52.9 | 3.2 |
| Fiber (%) | 4.7 | 4.8 |
| Ash (%) | 6.1 | 3 |
| Moisture (%) | <10 | <10 |
| Caloric profile (kcal/g) | 4.07 | 7.24 |
| Cystine (g/kg) | 2.8 | 0.3 |
| Methionine (g/kg) | 7 | 2.2 |

## REFERENCES

Alborzinia, Hamed, Andrés F. Flórez, Sina Kreth, Lena M. Brückner, Umut Yildiz, Moritz Gartlgruber, Dorett I. Odoni, Gernot Poschet, Karolina Garbowicz, Chunxuan Shao, Corinna Klein, Jasmin Meier, Petra Zeisberger, Michal Nadler-Holly, Matthias Ziehm, Franziska Paul, Jürgen Burhenne, Emma Bell, Marjan Shaikhkarami, Roberto Würth, Sabine A. Stainczyk, Elisa M. Wecht, Jochen Kreth, Michael Büttner, Naveed Ishaque, Matthias Schlesner, Barbara Nicke, Carlo Stresemann, María Llamazares-Prada, Jan H. Reiling, Matthias Fischer, Ido Amit, Matthias Selbach, Carl Herrmann, Stefan Wölfl, Kai Oliver Henrich, Thomas Höfer, Andreas Trumpp, and Frank Westermann. 2022. “MYCN Mediates Cysteine Addiction and Sensitizes Neuroblastoma to Ferroptosis.” Nature Cancer 3(4):471–85.

Anderson, Nicole M., Patrick Mucka, Joseph G. Kern, and Hui Feng. 2018. “The Emerging Role and Targetability of the TCA Cycle in Cancer Metabolism.” Protein and Cell 9(2):216–37.

Arbour, Kathryn C., Laura Mezquita, Niamh Long, Hira Rizvi, Edouard Auclin, Andy Ni, Gala Martínez-Bernal, Roberto Ferrara, W. Victoria Lai, Lizza E. L. Hendriks, Joshua K. Sabari, Caroline Caramella, Andrew J. Plodkowski, Darragh Halpenny, Jamie E. Chaft, David Planchard, Gregory J. Riely, Benjamin Besse, and Matthew D. Hellmann. 2018. “Impact of Baseline Steroids on Efficacy of Programmed Cell Death-1 and Programmed Death-Ligand 1 Blockade in Patients with Non-Small-Cell Lung Cancer.” Journal of Clinical Oncology 36(28):2872–78.

Badgley, Michael A., Daniel M. Kremer, H. Carlo Maurer, Kathleen E. DelGiorno, Ho Joon Lee, Vinee Purohit, Irina R. Sagalovskiy, Alice Ma, Jonathan Kapilian, Christina E. M. Firl,

Amanda R. Decker, Steve A. Sastra, Carmine F. Palermo, Leonardo R. Andrade, Peter Sajjakulnukit, Li Zhang, Zachary P. Tolstyka, Tal Hirschhorn, Candice Lamb, Tong Liu, Wei Gu, E. Scott Seeley, Everett Stone, George Georgiou, Uri Manor, Alina Iuga, Geoffrey M. Wahl, Brent R. Stockwell, Costas A. Lyssiotis, and Kenneth P. Olive. 2020. “Cysteine Depletion Induces Pancreatic Tumor Ferroptosis in Mice.” Science 368(6486):85–89.

Betancourt, Angelica, Sílvia Busquets, Marta Ponce, Míriam Toledo, Joan Guàrdia-Olmos, Maribel Peró-Cebollero, Francisco J. López-Soriano, and Josep M. Argilés. 2019. “The Animal Cachexia Score (ACASCO).” Animal Models and Experimental Medicine 2(3):201–9.

Bethin, Kathleen E., Sherri K. Vogt, and Louis J. Muglia. 2000. “Interleukin-6 Is an Essential, Corticotropin-Releasing Hormone-Independent Stimulator of the Adrenal Axis during Immune System Activation.” Proceedings of the National Academy of Sciences of the United States of America 97(16):9317–22.

Bhandarkar, Nikhil S., Rotem Lahav, Nitzan Maixner, Yulia Haim, G. William Wong, Assaf Rudich, and Uri Yoel. 2021. “Adaptation of Fuel Selection to Acute Decrease in Voluntary Energy Expenditure Is Governed by Dietary Macronutrient Composition in Mice.” Physiological Reports 9(18): 1–8.

Bratland, Eirik, Beate Skinningsrud, Dag E. Undlien, Edna Mozes, and Eystein S. Husebye. 2009. “T Cell Responses to Steroid Cytochrome P450 21-Hydroxylase in Patients with Autoimmune Primary Adrenal Insufficiency.” Journal of Clinical Endocrinology and Metabolism 94(12):5117–24.

Corbello Pereira, Sandra Regina, Elaine Darronqui, Jorgete Constantin, Mário Henrique Da Rocha Alves Da Silva, Nair Seiko Yamamoto, and Adelar Bracht. 2004. “The Urea Cycle and Related Pathways in the Liver of Walker-256 Tumor-Bearing Rats.” Biochimica et Biophysica Acta - Molecular Basis of Disease 1688(3):187–96.

Dello, Simon A. W. G., Evelien P. J. G. Neis, Mechteld C. de Jong, Hans M. H. van Eijk, Cécile H. Kicken, Steven W. M. Olde Damink, and Cornelis H. C. Dejong. 2013. “Systematic Review of Ophthalmate as a Novel Biomarker of Hepatic Glutathione Depletion.” Clinical Nutrition 32(3):325–30.

Divertie, Gavin D., Michael D. Jensen, and John M. Miles. 1991. “Stimulation of Lipolysis in Humans by Physiological Hypercortisolemia.” 40(October).

Dixon, Scott J., Kathryn M. Lemberg, Michael R. Lamprecht, Rachid Skouta, Eleina M. Zaitsev, Caroline E. Gleason, Darpan N. Patel, Andras J. Bauer, Alexandra M. Cantley, Wan Seok Yang, Barclay Morrison III, and Brent R. Stockwell. 2012. “Ferroptosis: An Iron-Dependent Form of Non-Apoptotic Cell Death.” Cell 149(5):1060–72.

Dong, Yang, Rongfu Tu, Hudan Liu, and Guoliang Qing. 2020. “Regulation of Cancer Cell Metabolism: Oncogenic MYC in the Driver’s Seat.” Signal Transduction and Targeted Therapy 5(1).

Efimova, luliia, Elena Catanzaro, Louis Van Der Meeren, Victoria D. Turubanova, Hamida Hammad, Tatiana A. Mishchenko, Maria V. Vedunova, Carmela Fimognari, Claus Bachert, Frauke Coppieters, Steve Lefever, Andre G. Skirtach, Olga Krysko, and Dmitri V. Krysko. 2020. “Vaccination with Early Ferroptotic Cancer Cells Induces Efficient Antitumor Immunity.” Journal for ImmunoTherapy of Cancer 8(2):1–15.

Eggermont, Alexander M. M., Michal Kicinski, Christian U. Blank, Mario Mandala, Georgina V. Long, Victoria Atkinson, Stéphane Dalle, Andrew Haydon, Adnan Khattak, Matteo S. Carlino, Shahneen Sandhu, James Larkin, Susana Puig, Paolo A. Ascierto, Piotr Rutkowski, Dirk Schadendorf, Rutger Koornstra, Leonel Hernandez-Aya, Anna Maria Di Giacomo, Alfonsus J. M. Van Den Eertwegh, Jean Jacques Grob, Ralf Gutzmer, Rahima Jamal, Paul C. Lorigan, Clemens Krepler, Nageatte Ibrahim, Sandrine Marreaud, Alexander Van Akkooi, Caroline Robert, and Stefan Suciu. 2020. “Association between Immune-Related Adverse Events and Recurrence-Free Survival among Patients with Stage III Melanoma Randomized to Receive Pembrolizumab or Placebo: A Secondary Analysis of a Randomized Clinical Trial.” JAMA Oncology 6(4):519–27.

Esteves, Francisco, José Rueff, and Michel Kranendonk. 2021. “The Central Role of Cytochrome P450 in Xenobiotic Metabolism—A Brief Review on a Fascinating Enzyme Family.” Journal ofXenobiotics 11(3):94–114.

Farkas, Jerneja, Stephan von Haehling, Kamyar Kalantar-Zadeh, John E. Morley, Stefan D. Anker, and Mitja Lainscak. 2013. “Cachexia as a Major Public Health Problem: Frequent, Costly, and Deadly.” Journal of Cachexia, Sarcopenia and Muscle 4(3):173–78.

Fearon, Kenneth, Florian Strasser, Stefan D. Anker, Ingvar Bosaeus, Eduardo Bruera, Robin L. Fainsinger, Aminah Jatoi, Charles Loprinzi, Neil MacDonald, Giovanni Mantovani, Mellar Davis, Maurizio Muscaritoli, Faith Ottery, Lukas Radbruch, Paula Ravasco, Declan Walsh, Andrew Wilcock, Stein Kaasa, and Vickie E. Baracos. 2011. “Definition and Classification of Cancer Cachexia: An International Consensus.” The Lancet Oncology 12(5):489–95.

Flint, Thomas R., Tobias Janowitz, Claire M. Connell, Edward W. Roberts, Alice E. Denton, Anthony P. Coll, Duncan I. Jodrell, and Douglas T. Fearon. 2016. “Tumor-Induced IL-6 Reprograms Host Metabolism to Suppress Anti-Tumor Immunity.” Cell Metabolism 24(5):672–84.

Fontana, Mario, Donatella Amendola, Emanuela Orsini, Alberto Boffi, and Laura Pecci. 2005. “Oxidation of Hypotaurine and Cysteine Sulphinic Acid by Peroxynitrite.” Biochemical Journal 389(l):233–40.

Fucà, Giovanni, Giulia Galli, Marta Poggi, Giuseppe Lo Russo, Claudia Proto, Martina Imbimbo, Roberto Ferrara, Nicoletta Zilembo, Monica Ganzinelli, Antonio Sica, Valter Torri, Mario Paolo Colombo, Claudio Vernieri, Andrea Balsari, Filippo De Braud, Marina Chiara Garassino, and Diego Signorelli. 2019. “Modulation of Peripheral Blood Immune Cells by Early Use of Steroids and Its Association with Clinical Outcomes in Patients with Metastatic Non-Small Cell Lung Cancer Treated with Immune Checkpoint Inhibitors.” ESMO Open 4(1):1—8.

Goldstein, Bernard D. 1975. “Mutagenicity of Malonaldehyde, a Decomposition Product of Peroxidized Polyunsaturated Fatty Acids.” Science 191(12).

Goncalves, Marcus D., Seo Kyoung Hwang, Chantal Pauli, Charles J. Murphy, Zhe Cheng, Benjamin D. Hopkins, David Wu, Ryan M. Loughran, Brooke M. Emerling, Guoan Zhang, Douglas T. Fearon, and Lewis C. Cantley. 2018. “Fenofibrate Prevents Skeletal Muscle Loss in Mice with Lung Cancer.” Proceedings of the National Academy of Sciences of the United States of America 115(4):E743–52.

de Groot, Stefanie, Rieneke T. Lugtenberg, Danielle Cohen, Marij J. P. Welters, llina Ehsan, Maaike P. G. Vreeswijk, Vincent T. H. B. M. Smit, Hiltje de Graaf, Joan B. Heijns, Johanneke E. A. Portielje, Agnes J. van de Wouw, Alex L. T. Imholz, Lonneke W. Kessels, Suzan Vrijaldenhoven, Arnold Baars, Elma Meershoek Klein Kranenbarg, Marjolijn Duijm de Carpentier, Hein Putter, Jacobus J. M. van der Hoeven, Johan W. R. Nortier, Valter D. Longo, Hanno Pijl, Judith R. Kroep, Hiltje de Graaf, Joan B. Heijns, Johanneke E. A. Portielje, Agnes J. van de Wouw, Alex L. T. Imholz, Lonneke W. Kessels, Suzan Vrijaldenhoven, Arnold Baars, Emine Göker, Anke J. M. Pas, Aafke H. Honkoop, A. Elise van Leeuwen-Stok, and Judith R. Kroep. 2020. “Fasting Mimicking Diet as an Adjunct to Neoadjuvant Chemotherapy for Breast Cancer in the Multicentre Randomized Phase 2 DIRECT Trial.” Nature Communications 11(1): 1—9.

von Haehling, Stephan and Stefan D. Anker. 2014. “Prevalence, Incidence and Clinical Impact of Cachexia: Facts and Numbers—Update 2014.” Journal of Cachexia, Sarcopenia and Muscle 5(4):261–63.

Haines, Ryan W., Parjam Zolfaghari, Yize Wan, Rupert M. Pearse, Zudin Puthucheary, and John R. Prowle. 2019. “Elevated Urea-to-Creatinine Ratio Provides a Biochemical Signature of Muscle Catabolism and Persistent Critical Illness after Major Trauma.” Intensive Care Medicine 45(12):1718–31.

Henzen, Christoph, Alex Suter, Erika Lerch, Ruth Urbinelli, Xaver H. Schorno, and Verena A. Briner. 2000. “Suppression and Recovery of Adrenal Response after Short-Term, High-Dose Glucocorticoid Treatment.” Lancet 355(9203):542–45.

Hoberman, H. D. 1950. “Endocrine Regulation of Amino Acid Protein Metabolism during Fasting.” The Yale Journal of Biology and Medicine 22(4):341–67.

Hopkins, Benjamin D., Chantal Pauli, Du Xing, Diana G. Wang, Xiang Li, David Wu, Solomon C. Amadiume, Marcus D. Goncalves, Cindy Hodakoski, Mark R. Lundquist, Rohan Bareja, Yan Ma, Emily M. Harris, Andrea Sboner, Himisha Beltran, Mark A. Rubin, Siddhartha Mukherjee, and Lewis C. Cantley. 2018. “Suppression of Insulin Feedback Enhances the Efficacy of PI3K Inhibitors.” Nature 560(7719):499–503.

Hsu, Jer Yuan, Suzanne Crawley, Michael Chen, Dina A. Ayupova, Darrin A. Lindhout, Jared Higbee, Alan Kutach, William Joo, Zhengyu Gao, Diana Fu, Carmen To, Kalyani Mondal, Betty Li, Avantika Kekatpure, Marilyn Wang, Teresa Laird, Geoffrey Horner, Jackie Chan, Michele Mcentee, Manuel Lopez, Damodharan Lakshminarasimhan, Andre White, Sheng Ping Wang, Jun Yao, Junming Yie, Hugo Matern, Mark Solloway, Raj Haldankar, Thomas Parsons, Jie Tang, Wenyan D. Shen, Yu Alice Chen, Hui Tian, and Bernard B. Allan. 2017. “Non-Homeostatic Body Weight Regulation through a Brainstem-Restricted Receptor for GDF15.” Nature 550(7675):255–59.

Hursting, Stephen D. and Nathan A. Berger. 2010. “Energy Balance, Host-Related Factors, and Cancer Progression.” Journal of Clinical Oncology 28(26):4058–65.

Janowitz, Tobias. 2018. “Cancer: The Tumor-Driven Disease of the Host.” Cell Metabolism 28(l):5–6.

Jansen, Natalie and Harald Walach. 2016. “The Development of Tumours under a Ketogenic Diet in Association with the Novel Tumour Marker TKTL1: A Case Series in General Practice.” Oncology Letters ll(l):584–92.

Lien, Evan C., Anna M. Westermark, Yin Zhang, Chen Yuan, Zhaoqi Li, Allison N. Lau, Kiera M. Sapp, Brian M. Wolpin, and Matthew G. Vander Heiden. 2021. “Low Glycaemic Diets Alter Lipid Metabolism to Influence Tumour Growth.” Nature 599(7884):302–7.

Little, Clive and Peter J. O’Brien. 1968. “An Intracellular GSH-Peroxidase with a Lipid Peroxide Substrate.” Biochemical and Biophysical Research Communications 31(2):145–50.

Liu, Dora, Alexandra Ahmet, Leanne Ward, Preetha Krishnamoorthy, Efrem D. Mandelcorn, Richard Leigh, Jacques P. Brown, Albert Cohen, and Harold Kim. 2013. “A Practical Guide to the Monitoring and Management of the Complications of Systemic Corticosteroid Therapy.” Allergy, Asthma and Clinical Immunology 9(l):l-25.

Lu, Yuxiong, Qing Yang, Yubin Su, Yin Ji, Guobang Li, Xianzhi Yang, Liyan Xu, Zhaoliang Lu, Jiajun Dong, Yi Wu, Jin Xin Bei, Chaoyun Pan, Xiaoqiong Gu, and Bo Li. 2021. “MYCN Mediates TFRC-Dependent Ferroptosis and Reveals Vulnerabilities in Neuroblastoma.” Cell Death and Disease 12(6).

Maddocks, Oliver D. K., Dimitris Athineos, Eric C. Cheung, Pearl Lee, Tong Zhang, Niels J. F. Van Den Broek, Gillian M. Mackay, Christiaan F. Labuschagne, David Gay, Flore Kruiswijk, Julianna Blagih, David F. Vincent, Kirsteen J. Campbell, Fatih Ceteci, Owen J. Sansom, Karen Blyth, and Karen H. Vousden. 2017. “Modulating the Therapeutic Response of Tumours to Dietary Serine and Glycine Starvation.” Nature 544(7650):372–76.

Mager, Donald E., Sheren X. Lin, Robert A. Blum, Christian D. Lates, and William J. Jusko. 2003. “Dose Equivalency Evaluation of Major Corticosteroids: Pharmacokinetics and Cell Trafficking and Cortisol Dynamics.” Journal of Clinical Pharmacology 43(**ll**):1216-27.

Massey, Karen A. and Anna Nicolaou. 2011. “Lipidomics of Polyunsaturated-Fatty-Acid-Derived Oxygenated Metabolites.” Biochemical Society Transactions 39(5):1240–46.

Massucci, Maria, Francesca Di Fabio, Fabiola L. Rojas Llimpe, and Andrea Ardizzoni. 2020. “A Case of Response to Immunotherapy in a Patient with MSI Metastatic Colorectal Cancer and Autoimmune Disease in Steroid Therapy.” Journal of Immunotherapy 43(5):153–55.

Menzies, Alexander M., D. B. Johnson, S. Ramanujam, V. G. Atkinson, A. N. M. Wong, J. J. Park, J. L. McQuade, A. N. Shoushtari, K. K. Tsai, Z. Eroglu, O. Klein, J. C. Hassel, J. A. Sosman, A. Guminski, R. J. Sullivan, A. Ribas, M. S. Carlino, M. A. Davies, S. K. Sandhu, and G. V. Long. 2017. “Anti-PD-1 Therapy in Patients with Advanced Melanoma and Preexisting Autoimmune Disorders or Major Toxicity with Ipilimumab.” Annals of Oncology 28(2):368–76.

Mishima, Eikan. 2021. “The E2F1-IREB2 Axis Regulates Neuronal Ferroptosis in Cerebral Ischemia.” Hypertension Research 1085–86.

Mulderrig, Lee, Juan I. Garaycoechea, Zewen K. Tuong, Christopher L. Millington, Felix A. Dingier, John R. Ferdinand, Liam Gaul, John A. Tadross, Mark J. Arends, Stephen O’Rahilly, Gerry P. Crossan, Menna R. Clatworthy, and Ketan J. Patel. 2021. “Aldehyde-Driven Transcriptional Stress Triggers an Anorexic DNA Damage Response.” Nature 600(7887):158–63.

Nair, Jagadeesan, Carlos E. Vaca, Ivana Velic, Marja Mutanen, Liisa M. Valsta, and Helmut Bartsch. 2019. “High Dietary W-6 Polyunsaturated Fatty Acids Drastically Increase the Formation of Etheno-DNA Base Adducts in White Blood Cells of Female Subjects.” Journal of Chemical Information and Modeling 53(9):1689–99.

Nakamura, Kentaro, Hidekazu Tonouchi, Akina Sasayama, and Kinya Ashida. 2018. “A Ketogenic Formula Prevents Tumor Progression and Cancer Cachexia by Attenuating Systemic Inflammation in Colon 26 Tumor-Bearing Mice.” Nutrients 10(2).

Otto, Christoph, Ulrike Kaemmerer, Bertram Illert, Bettina Muehling, Nadja Pfetzer, Rainer Wittig, Hans Ullrich Voelker, Arnulf Thiede, and Johannes F. Coy. 2008. “Growth of Human Gastric Cancer Cells in Nude Mice Is Delayed by a Ketogenic Diet Supplemented with Omega-3 Fatty Acids and Medium-Chain Triglycerides.” BMC Cancer 8:1–12.

Pannala, Venkat R., Jason N. Bazil, Amadou K. S. Camara, and Ranjan K. Dash. 2013. “A Biophysically-Based Mathematical Model for the Catalytic Mechanism of Glutathione Reductase.” Free Radical Biology and Medicine 65.

Patel, Satish, Anna Alvarez-Guaita, Audrey Melvin, Debra Rimmington, Alessia Dattilo, Emily L. Miedzybrodzka, Irene Cimino, Anne Catherine Maurin, Geoffrey P. Roberts, Claire L. Meek, Samuel Virtue, Lauren M. Sparks, Stephanie A. Parsons, Leanne M. Redman, George A. Bray, Alice P. Liou, Rachel M. Woods, Sion A. Parry, Per B. Jeppesen, Anders J. Koines, Heather P. Harding, David Ron, Antonio Vidal-Puig, Frank Reimann, Fiona M. Gribble, Carl J. Hulston, I. Sadaf Farooqi, Pierre Fafournoux, Steven R. Smith, Jorgen Jensen, Danna Breen, Zhidan Wu, Bei B. Zhang, Anthony P. Coll, David B. Savage, and Stephen O’Rahilly. 2019. “GDF15 Provides an Endocrine Signal of Nutritional Stress in Mice and Humans.” Cell Metabolism 29(3):707–718.e8.

Pissios, Pavlos, Shangyu Hong, Adam Richard Kennedy, Deepthi Prasad, Fen Fen Liu, and Eleftheria Maratos-Flier. 2013. “Methionine and Choline Regulate the Metabolic Phenotype of a Ketogenic Diet.” Molecular Metabolism 2(3):306–13.

Rui, Liangyou. 2014. “Energy Metabolism in the Liver.” Compr Physiol. 4(1):177–97.

Salas, M. A., S. W. Evans, M. J. Levell, and J. T. Whicher. 1990. “Interleukin-6 and ACTH Act Synergistically to Stimulate the Release of Corticosterone from Adrenal Gland Cells.” Clinical and Experimental Immunology 79(3):470–73.

Schwartz, Kenneth, Howard T. Chang, Michele Nikolai, Joseph Pernicone, Sherman Rhee, Karl Olson, Peter C. Kurniali, Norman G. Hord, and Mary Noel. 2015. “Treatment of Glioma Patients with Ketogenic Diets: Report of Two Cases Treated with an IRB-Approved Energy- Restricted Ketogenic Diet Protocol and Review of the Literature.” Cancer & Metabolism 3(l):l-10.

Scuto, Maria, Angela Trovato Salinaro, Sergio Modafferi, Alessandra Polimeni, Tilman Pfeffer, Tim Weigand, Vittorio Calabrese, Claus Peter Schmitt, and Verena Peters. 2020. “Carnosine Activates Cellular Stress Response in Podocytes and Reduces Glycative and Lipoperoxidative Stress.” Biomedicines 8(6):1–14.

Seyfried, T. N., T. M. Sanderson, M. M. El-Abbadi, R. McGowan, and P. Mukherjee. 2003. “Role of Glucose and Ketone Bodies in the Metabolic Control of Experimental Brain Cancer.” British Journal of Cancer 89(7):1375–82.

Ursini, F., M. Maiorino, M. Valente, L. Ferri, and C. Gregolin. 1982. “Purification from Pig Liver of a Protein Which Protects Liposomes and Biomembranes from Peroxidative Degradation and Exhibits Glutathione Peroxidase Activity on Phosphatidylcholine Hydroperoxides.” Biochimica et Biophysica Acta (BBA)/Lipids and Lipid Metabolism 710(2):197–211.

Vicanolo, Tommaso, Andres Hidalgo, and Jose M. Adrover. n.d. Measuring Circadian Neutrophil Infiltration in Tissues by Paired Whole-Mount Tissue Clearing and Flow Cytometry. Vol. 2482.

Warburg, Otto. 1925. “The Metabolism of Carcinoma Cells 1.” The Journal of Cancer Research 9(l):148–63.

Wing, S. S. and A. L. Goldberg. 1993. “Glucocorticoids Activate the ATP-Ubiquitin-Dependent Proteolytic System in Skeletal Muscle during Fasting.” American Journal of Physiology - Endocrinology and Metabolism 264(4 27-4).

Yang, Wan Seok, Rohitha Sriramaratnam, Matthew E. Welsch, Kenichi Shimada, Rachid Skouta, Vasanthi S. Viswanathan, Jaime H. Cheah, Paul A. Clemons, F. Shamji, Clary B. Clish, Lewis M. Brown, Albert W. Girotti, Virginia W. Cornish, Stuart L. Schreiber, Brent R. Stockwell, West Street, and New York. 2014. “Regulation of Ferroptotic Cancer Cell Death by GPX4.” Ceil 156(0):317–31.

Yin, Huiyong, Libin Xu, and Ned A. Porter. 2011. “Free Radical Lipid Peroxidation: Mechanisms and Analysis.” Chemical Reviews lll(10):5944–72.

Yuan, Lei, Jun Han, Qingyang Meng, Qiulei Xi, Qiulin Zhuang, Yi Jiang, Yusong Han, Bo Zhang, Jing Fang, and Guohao Wu. 2015. “Muscle-Specific E3 Ubiquitin Ligases Are Involved in Muscle Atrophy of Cancer Cachexia: An in Vitro and in Vivo Study.” Oncology Reports 33(5):2261–68.

Žarković, Miloš, Svetlana Ignjatović, Marijana Dajak, Jasmina Ćirić, Biljana Beleslin, Slavica Savić, Mirjana Stojković, Petar Bulat, and Božo Trbojević. 2008. “Cortisol Response to ACTH Stimulation Correlates with Blood Interleukin 6 Concentration in Healthy Humans.” European Journal of Endocrinology 159(5):649–52.

