## Supplementary material for "Ketogenic diet promotes tumor ferroptosis but induces relative corticosterone deficiency that accelerates cachexia": Key Resources Table

| REAGENT or RESOURCE | SOURCE | IDENTIFIER |
| --- | --- | --- |
| Antibodies | | |
| anti-human/mouse Myeloperoxidase (MPO) | R&D Systems | Cat. #AF3667; RRID:AB_2250866 |
| Donkey anti-Goat IgG (H+L) Alexa Fluor 633 | Thermo Fisher | Cat. #A-21082; RRID:AB_10562400 |
| anti-Ki67 | Thermo Fisher | Cat. #14-5698-82; RRID:AB_10854564 |
| anti-mouse CD45 Alexa Fluor 700 | BioLegend | Cat. #103127; RRID:AB_493714 |
| anti-mouse CD45 FITC | Thermo Fisher | Cat. #11-0451; RRID:AB_465049 |
| anti-mouse CD3ε APC/Cy7 | BioLegend | Cat. #100329; RRID:AB_1877171 |
| anti-mouse CD4 PerCP/Cyanine 5.5 | BioLegend | Cat. #100433; RRID:AB_893330 |
| anti-mouse CD8a Brilliant Violet 510TM | BioLegend | Cat. #100751; RRID:AB_2561389 |
| anti-mouse CD11b Brilliant Violet 605TM | BioLegend | Cat. #101257; RRID:AB_2565431 |
| anti-mouse Ly-6G/Ly-6C Alexa Fluor 700 | BioLegend | Cat. #108421; RRID:AB_493728 |
| anti-mouse CD69 FITC | BioLegend | Cat. #104505; RRID:AB_313108 |
| anti-mouse CD152 PE/Cy7 | BioLegend | Cat. #106313; RRID:AB_2564237 |
| anti-mouse CD274 Brilliant Violet 421TM | BioLegend | Cat. #124315; RRID:AB_10897097 |
| anti-mouse CD279 PE/Dazzle 594 | BioLegend | Cat. #109115; RRID:AB_2566547 |
| anti-mouse F4/80 FITC | BioLegend | Cat. #123107; RRID:AB_893500 |
| Rabbit anti-BAX mAb | Proteintech | Cat. #50599-2-Ig; RRID:AB_2061561 |
| Rabbit β-Actin mAb | Cell Signaling | Cat. #4967; RRID:AB_330288 |
| Rabbit Caspase-3 (D3R6Y) mAb | Cell Signaling | Cat. #14220; RRID:AB_2798429 |
| Goat Anti-Rabbit IgG H&L (HRP) | Abcam | Cat. # ab205718; RRID:AB_2819160 |
| Chemicals, peptides, and recombinant proteins | | |
| Dexamethasone 21-phosphate disodium salt | Sigma-Aldrich | Cat. #D1159 |
| N-Acetyl-L-cysteine | Sigma-Aldrich | Cat. #A9165 |
| Sodium Chloride (NaCl) 0.9% sterile saline | Thermo Fisher | Cat. #Z1377 |
| RSL3 ((1S,3R)-RSL3) | MedChemExpress | Cat. #HY-100218A |
| Dimethyl Sulfoxide (DMSO) | Cell Signaling | Cat. #12611S |
| Polyethylene glycol 300 (PEG300) | Selleck Chemicals | Cat. #S6704 |
| ACTH | ProSpec | Cat. #HOR-279 |
| 2x Lysis buffer | RayBiotech | Cat. #AA-LYS-16ml |
| Protease Inhibitor Cocktail | RayBiotech | Cat. #AA-PI |
| Phosphatase Inhibitor Cocktail Set I | RayBiotech | Cat. #AA-PHI-I |
| Formalin | Sigma-Aldrich | Cat. # HT501128-4L |
| Propylene Glycol | Sigma-Aldrich | Cat. #P4347 |
| Oil Red O | Sigma-Aldrich | Cat. #O0625-25G |
| Hematoxylin and Eosin Kit | Abcam | Cat. #ab245880 |
| Methanol | VWR | Cat. #BDH1135-4LP |
| Benzyl alcohol | Sigma-Aldrich | Cat. #108006-500ML |
| Benzyl benzoate | Sigma-Aldrich | Cat. #B6630-500ML |
| Paraformaldehyde | Thermo Fisher | Cat. #50-980-495 |
| Fetal Bovine Serum (FBS) | Thermo Fisher | Cat. #10-438-026 |
| RPMI-640 | Thermo Fisher | Cat. #11-875-093 |
| Penicillin-Streptomycin | Thermo Fisher | Cat. #15-140-122 |
| Trypsin-EDTA (0.5%) | Thermo Fisher | Cat. #15400054 |
| DMEM:F12 | ATCC | Cat. #30-2006 |
| Nu-Serum I | Corning | Cat. #355100 |
| ITS+ Premix | Corning | Cat. #354352 |
| 4-hydroxynonenal (4-HNE) | Cayman Chemical | Cat. #32100 |
| 4-hydroxyhexenal (4-HHE) | Cayman Chemical | Cat. #32060 |
| Malondialdehyde (MDA) | Sigma-Aldrich | Cat. #63287-1G-F |
| Trypan blue | Thermo Fisher | Cat. #15250061 |
| Trichloroacetic acid | Sigma-Aldrich | Cat. #T9159-100G |
| Sulforhodamine B (SRB) | Sigma-Aldrich | Cat. #S1402-5G |
| Acetic acid | Sigma-Aldrich | Cat. #A6283 |
| Collagenase I | Sigma-Aldrich | Cat. #SCR103 |
| DNase I | Sigma-Aldrich | Cat. #04716728001 |
| Percoll | Sigma-Aldrich | Cat. #GE17-0891-01 |
| RBC lysis buffer | Thermo Fisher | Cat. #A1049201 |
| RIPA buffer | Thermo Fisher | Cat. #89901 |
| Dithiothreitol (DTT) | Thermo Fisher | Cat. #A39225 |
| Blotting-Grade Blocker (dry milk) | Bio-Rad | Cat. #1706404 |
| ECL | Thermo Fisher | Cat. #32106 |
| QIAzol Lysis Reagent | Qiagen | Cat. #79396 |
| Critical commercial assays | | |
| Corticosterone ELISA | IBL | Cat. #RE52211 |
| IL-6 Quantikine ELISA | R&D | Cat. #M6000B |
| 4-Hydroxynonenal ELISA | Novus Biologicals | Cat. #NBP2-66364 |
| Pregnenolone ELISA | Novus Biologicals | Cat. #NBP2-68102 |
| Progesterone ELISA | Novus Biologicals | Cat. #NBP2-60125 |
| Leptin ELISA | Thermo Fisher | Cat. #KMC2281 |
| ACTH ELISA | Abcam | Cat. #ab263880 |
| NADP/NADPH-Glo™ Bioluminescent Assay | Promega | Cat. #G9081 |
| Iron assay | Abcam | Cat. #ab83366 |
| Lipid Peroxidation (4-HNE) Assay | Abcam | Cat. #ab238538 |
| Cortisol Competitive ELISA | Thermo Fisher | Cat. #EIAHCOR |
| RNeasy Lipid Tissue Mini Kit | Qiagen | Cat. #74804 |
| TaqMan™ RNA-to-CT™ 1-Step Kit | Thermo Fisher | Cat. #4392653 |
| Experimental models: Cell lines | | |
| C26 murine colorectal cancer |  | RRID:CVCL_3925 |
| H295R human adrenocortical cancer | ATCC | Cat. #CRL-2128; RRID:CVCL_0458 |
| Experimental models: Organisms/strains |  |  |
| BALB/c | Charles River | Cat. #028 |
| KPC |  |  |
| Software and algorithms | | |
| FlowJo X (Tree Star) | FlowJo LLC | <https://www.flowjo.com> |
| CLAMS/Oxymax | Columbus Instruments | Comprehensive Lab Animal Monitoring System (CLAMS) |
| GraphPad Prism 9 | GraphPad Software | <https://www.graphpad.com> |
| ImageJ | NIH | <https://imagej.nih.gov/ij/> |
| Other | | |
| Standard chow diet (PicoLab® Rodent Diet) | LabDiet | Cat. #5053 |
| Ketogenic diet (KD) | Bio-Serv | Cat. #F3666 |
| 35mm Dishes with 1.5 Coverslip and 10 mm Glass Diameter | MatTek | Cat. #P35G-1.5-10-C |
| Nitrocellulose membranes | Thermo Fisher | Cat. #88025 |
